# Plant-based production of diverse human milk oligosaccharides

**DOI:** 10.1101/2023.09.18.558286

**Authors:** Collin R. Barnum, Bruna Paviani, Garret Couture, Chad Masarweh, Ye Chen, Yu-Ping Huang, David A. Mills, Carlito B. Lebrilla, Daniela Barile, Minliang Yang, Patrick M. Shih

## Abstract

Human milk oligosaccharides (HMOs) are a diverse class of carbohydrates that aid in the health and development of infants. The vast health benefits of HMOs have made them a commercial target for microbial production; however, producing the ∼130 structurally diverse HMOs at scale has proven difficult. Here, we produce a vast diversity of HMOs by leveraging the robust carbohydrate anabolism of plants. This diversity includes high value HMOs, such as lacto-N-fucopentaose I, that have not yet been commercially produced using state-of-the-art microbial fermentative processes. HMOs produced in transgenic plants provided strong bifidogenic properties, indicating their ability to serve as a prebiotic supplement. Technoeconomic analyses demonstrate that producing HMOs in plants provides a path to the large-scale production of specific HMOs at lower prices than microbial production platforms. Our work demonstrates the promise in leveraging plants for the cheap and sustainable production of HMOs.

## Introduction

Human milk is a complete and comprehensive food evolved to nourish and protect infants. A key component to the distinct bioactive properties of human milk is the presence of a wide diversity of human milk oligosaccharides (HMOs) that are well documented in establishing the nascent gut microbiota of infants to prevent diseases and ensure healthy development^1–4^. While 75% of infants are supplemented with or exclusively fed infant formula in the first six months of life, current infant formulas are either devoid of HMOs or only contain one to two of the ∼130 HMOs found in human milk, limiting the health outcomes of formula-fed infants^3,5^. In addition to their use for infant health, HMOs are being studied for their beneficial roles in adult health as a prebiotic to improve intestinal barrier function, lower gastrointestinal inflammation, and treat irritable bowel diseases^6–10^; however, the study of HMO benefits in adults has been limited to a small subset of HMOs. Currently, commercial HMO production relies on microbial fermentation, but, to date, microbial fermentation is only able to commercially produce two to five simple HMOs of the ∼130 HMOs found in human milk at scale^11,12^. While five simple HMOs constitute a large portion of HMO mass in human milk, diverse HMOs with a range of linkages and degrees of polymerization enable the growth of beneficial gut microbes that have preferences for specific HMOs^13,14^. Thus, there is a need to develop novel platforms to produce a wider diversity of HMOs found in human milk that can improve the bioactive properties of infant formula to improve the health of infants and adults worldwide.

The combinatorial nature of glycosidic linkages, nucleotide sugar donors, and oligosaccharide acceptor molecules enables the large diversity of HMOs found in human milk^15^. HMOs are composed of five distinct sugars – D-glucose (Glc), D-galactose (Gal), *N*-acetylglucosamine (GlcNAc), L-fucose (Fuc), and *N*-acetylneuraminic acid (Neu5Ac) – connected via various glycosidic linkages to generate a diverse range of molecular structures **(Fig. 1A)**. HMO biosynthesis begins with the production of lactose that can be decorated with fucose or Neu5Ac to form a variety of tri- and tetrasaccharides. Lactose can also be extended by glycosyltransferases through the addition of a GlcNAc-ß-1,3 or GlcNAc-ß-1,4 to form unbranched HMOs. Unbranched HMOs can be further extended by glycosyltransferases through the addition of a GlcNAc-ß-1,6 to produce branched HMOs **(Fig. 1B)**. GlcNAc can undergo subsequent addition of Gal-ß-1,3 or Gal-ß-1,4 to form type I and type II HMOs, respectively **(Fig. 1B)**. While HMOs can consist of all five monosaccharides, they are generally classified in three broad HMO groups based on their composition: 1) Neutral HMOs contain Glc, Gal, and GlcNAc, 2) Fucosylated HMOs contain a neutral core with one or more Fuc additions, and 3) Acidic HMOs contain a neutral core with one or more additions of Neu5Ac. Due to the need for high amounts of nucleotide sugars and glycosylation potential in HMO biosynthesis, a suitable host must have robust sugar metabolism capable of managing the metabolic burden of HMO production.

**Figure 1.**
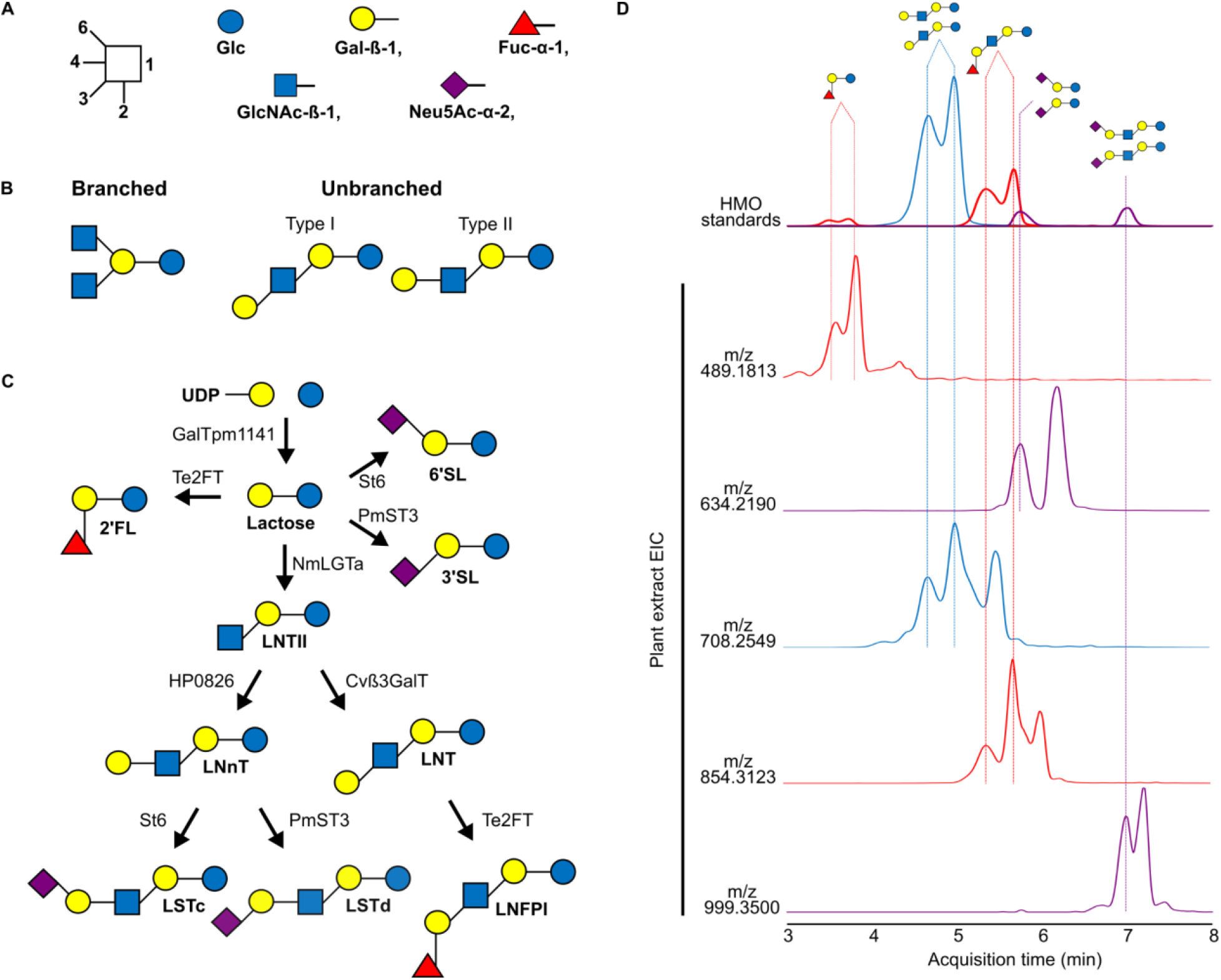
Production of all three HMO classes *in planta*. A) HMOs are composed of glucose (Glc), galactose (Gal), fucose (Fuc), *N*-acetylglucosamine (GlcNAc), and/or *N*-acetylneuraminic acid (Neu5Ac) connected via Gal-ß-1,3/4-, GlcNAc-ß-1,3/6-, Fuc-α-1,2/3/4, or Neu5Ac-α-2,3/6 glycosidic linkages. B) HMOs can be divided into branched, unbranched, type I and/or type II HMOs. C) HMO biosynthetic pathways used in this study for HMO production *in planta*. D) Extracted ion chromatograms displaying the identification of 2’FL, 3’SL, 6’SL, LNFPI, LSTa, LSTc, and LNT/LNnT in extracts of individual plant leaves using LC-MS/MS (Q Exactive, Thermo). Red, blue and purple coloring denote fucosylated, neutral and acidic HMOs, respectively. Some additional peaks are present due to in-source fragmentation of larger oligosaccharides with no available standards.

Unlike many microbes used in commercial fermentation, plants have evolved to create a wide range of glycans that encompass a diversity of nucleotide sugars from photosynthetically fixed CO_2_. As masters of sugar anabolism, plants are able to create vast amounts of complex oligo- and polysaccharides^16^. This has led to commercial operations for the production of prebiotic oligosaccharides from plant biomass, such as ß-glucan, xylooligosaccharides, inulin or soy oligosaccharides^17–20^. Many of these products can either be purified or directly consumed as a food, providing an easy means of ingestion. Additionally, plants can be grown in open fields, requiring minimal inputs, limiting the need for expensive substrates and axenic conditions^21^. The robust sugar metabolism of plants and ability to be grown at agricultural scales make plants an ideal platform for the large scale production of HMOs.

Due to the unique advantages of plants as a platform for carbohydrate production, we tested their ability to produce a range of HMOs using both transient and stable expression in *Nicotiana benthamiana*. Here, we report the first *in planta* production of neutral, fucosylated, and acidic HMOs, including several HMOs never before produced in a heterologous host, showcasing the intrinsic advantages of a plant-based production platform. Furthermore, we show that plant-produced HMOs provide selective growth of key bifidobacteria, suggesting their potential prebiotic efficacy. Finally, we assess the economic viability of HMO production *in planta* compared to current microbial platforms.

## Results

### Production of all three HMO classes in planta

Production of HMOs requires the expression of glycosyltransferases capable of creating specific glycosidic linkages. While HMO biosynthesis in humans takes place in the Golgi via a largely unknown pathway^22^, various microbial enzymes are capable of generating HMOs in the cytosol. To produce HMOs in plants, we localized bacterial HMO biosynthetic enzymes to the cytosol for the production of neutral, fucosylated, and acidic HMOs **(Fig. 1C)**. We tested these pathways through transient expression in *Nicotiana benthamiana*. Transient expression permits relatively high throughput screening of biosynthetic pathways *in planta* by injecting strains of *Agrobacterium tumefaciens* into plant leaves, allowing the *Agrobacterium* to insert genes for HMO biosynthetic genes into the plant cell^23^. Leaves transiently expressing HMO biosynthetic pathways were subjected to liquid-liquid extraction, C18 solid-phase extraction (SPE) and porous graphitic carbon (PGC) SPE before characterization by mass spectrometry **(Supplementary Fig. 1)**.

Neutral HMOs function as the core scaffolds of other more complex HMOs (i.e., fucosylated and acidic); thus, we first targeted the type I and type II neutral HMO core structures, lacto-*N*-tetraose (LNT) and lacto-*N*-neotetraose (LNnT). Expression of a neutral HMO biosynthetic pathway using two ß-1,4-galactosyltransferases (*GalTPM1141*^24^, *Hp0826*^25^), one ß-1,3-galactosyltransferase (*Cvß3GalT*^26^), and one ß-1,3-*N*-acetylglucosaminyltransferease (*NmLgtA*^27^) resulted in the production of lactose and various neutral HMOs with degrees of polymerization ranging from three to seven. Notably, *N. benthamiana* transiently expressing this pathway produced the tetrasaccharides LNT (*m/z* 708.2559) and LNnT (*m/z* 708.2559), which represent major type I and type II HMOs in human milk, respectively^28^ **(Fig. 1D, Supplementary Table 1)**. Additionally, we identified the production of larger neutral oligosaccharides with varying degrees of polymerization using MS/MS fragmentation to determine the number of hexose and HexNAc sugars **(Supplementary Table 1)**. This included multiple neutral isomers of pentasaccharides and heptasaccharides **(Supplementary Table 1)**. Our findings demonstrate that plants have the ability to produce previously inaccessible oligosaccharides with various degrees of polymerization that may exhibit novel bioactivity.

Following the success of generating type I and type II neutral HMOs, we examined the ability of plants to decorate neutral HMOs with fucose, as fucosylated HMOs are the most abundant class of HMO in human milk^14^. We transiently expressed an α-1,2-fucosyltransferase (*Te2FT*^29^) alongside the neutral HMO biosynthetic pathway to produce the most abundant fucosylated HMOs in human milk: 2’-fucosyllactose (2’FL) (*m/z* 489.1819) and lacto-*N*-fucopentaose I (LNFPI) (*m/z* 854.3136) **(Fig. 1D, Supplementary Table 1)**. Additionally, several fucosylated hexasaccharide isomers were identified by *m/z* and MS/MS fragmentation **(Supplementary Table 1)**. While the structure of each isomer could not be determined, each is composed of four hexoses, one HexNAc, and one deoxyhexose, indicating that either LNFPI can be further decorated with hexose sugars or pentasaccharide neutral HMOs can be decorated with additional fucose.

While neutral and fucosylated HMOs represent a majority of HMOs in human milk, acidic HMOs constitute the last major class found in mammalian milks which provide unique bioactivities due to the presence of *N*-acetylneuraminic acid^30^. Plants do not natively produce the donor molecule for production of acidic HMOs, CMP-Neu5Ac. To produce acidic HMOs, we simultaneously expressed the neutral HMO biosynthetic pathway, sialyltransferases, and a mammalian pathway for the production of CMP-Neu5Ac **(Supplementary Fig. 2)**^31^. Expression of an α-2,6-sialyltransferase (*St6*^32^) alongside the neutral HMO biosynthetic pathway produced the acidic trisaccharide, 6’-sialyllactose (6’SL) (*m/z* 634.2191), of type II acidic HMO, sialyllacto-*N*-neotetraose c (LSTc) (*m/z* 999.3505) **(Fig. 1D, Supplementary Table 1)**. Expression of an α-2,3-sialyltransferase (*PmST3*^33^) with the neutral HMO biosynthetic pathway produced a myriad of acidic HMOs, such as the acidic trisaccharide, 3’-sialyllactose (3’SL) (*m/z* 634.2187), and the acidic pentasaccharide, LSTd (*m/z* 999.3510) **(Fig. 1D, Supplementary Table. 1)**. These findings mark the first heterologous production of LST isomers *in vivo.* Additionally, six isomers of acidic hexasaccharides were identified using *m/z* and MS/MS fragmentation. Each isomer was composed of four hexoses, one HexNAc, and one Neu5Ac **(Supplementary Table 1)**. Together, these results display the ability of plants to produce all three classes of HMOs combinatorially or simultaneously **(Supplementary Fig. 3)**, including type I and type II structures, marking the greatest diversity of HMOs made in a single heterologous organism.

### Optimized production of complex fucosylated HMOs in N. benthamiana

Microbial production platforms suffer from an inability to produce HMOs with higher degrees of polymerization at large scales, leaving many larger, more complex HMOs understudied. LNFPI is a fucosylated pentasaccharide that is the second most abundant fucosylated HMO. Despite its high abundance in breast milk, LNFPI has remained recalcitrant to fermentative production in microbes, limiting efforts to study its potential health benefits. Therefore, we sought to optimize the production of LNFPI *in planta* by overexpressing the requisite nucleotide sugar biosynthetic pathways. We transiently expressed the biosynthetic pathway for LNFPI **(Fig. 2A)** alongside pathways for the production of UDP-galactose, UDP-*N*-acetylglucosamine (UDP-GlcNAc), and GDP-fucose and quantified LNFPI production **(Fig. 2B)**. Expression of the LNFPI pathway with the GDP-fucose pathway increased production of LNFPI by 32.9% (1075.03 ug/g dry weight) compared to the expression of the LNFPI pathway alone (808.91 ug/g dry weight), indicating that GDP-fucose is limiting in *N. benthamiana* **(Fig. 2C)**. Surprisingly, overexpression of the GDP-fucose pathway also resulted in the production of lactodifucotetraose (LDFT) (*m/z* 635.2394) and lacto-*N*-difuco-hexaose I (LNDFHI) (*m/z* 1000.3720) **(Supplementary Table 1)** despite not expressing an α-1,3- or α-1,4-fucosyltransferase, indicating the presence of a native plant fucosyltransferases capable of glycosylating HMOs. Overexpression of all other nucleotide sugar pathway combinations resulted in similar or lower levels of LNFPI production compared to expression of the LNFPI pathway alone.

**Figure 2.**
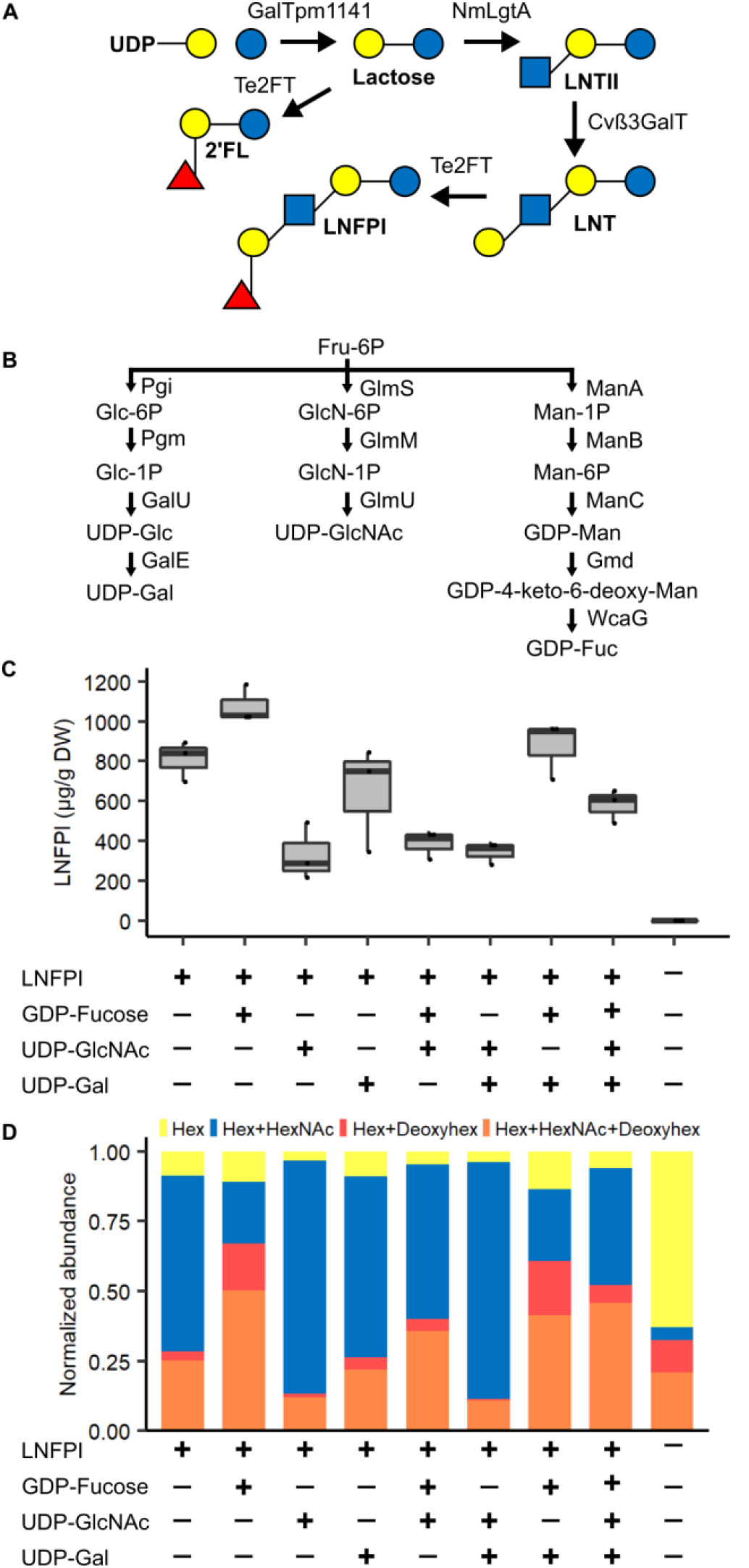
Manipulation of nucleotide sugar biosynthetic pathways modulates HMO profiles *in planta*. A) HMO biosynthetic pathway expressed for the production of LNFPI. B) Nucleotide sugar biosynthetic pathways expressed with LNFPI pathway. C) Quantification of LNFPI production through expression of LNFPI biosynthetic pathway alongside combinatorially expressed nucleotide sugar biosynthetic pathways using an internal calibration curve obtained with an Agilent 6530 Accurate-Mass Q-TOF MS. D) Effect of nucleotide sugar biosynthetic pathway overexpression on HMO profile produced using the LNFPI pathway. Composition based on Hexose, HexNAc, Deoxyhexose (Deoxyhex) composition determined using *m/z* and MS/MS fragmentation. Mass spectral analysis was performed on an Agilent Q-TOF MS.

Interestingly, the overall profile of oligosaccharides produced was altered by overexpressing nucleotide sugar biosynthetic pathways **(Fig. 2D)**. The number of hexose (Glc, Gal), HexNAc (GlcNAc), and deoxyhexose (Fuc) sugars in each oligosaccharide identified was determined through identification via *m/z* and MS/MS fragmentation and normalized using a LNFPI calibration curve. Overexpression of the UDP-GlcNAc and LNFPI pathways increased the relative amount of neutral oligosaccharides containing a hexose and HexNAc compared to expression of the LNFPI pathway alone **(Fig. 2D)**. Overexpression of the GDP-fucose and LNFPI pathways resulted in a shift in the overall oligosaccharide composition, favoring the production of oligosaccharides containing at least one deoxyhexose, indicating an increase in the level of fucosylated oligosaccharides **(Fig. 2D)**. These results demonstrate that tailoring the availability of nucleotide sugars enables control over the ratio of HMOs produced.

Because scaling HMO production in plants requires the growth of stably transformed plants, we developed transgenic lines of *N. benthamiana* expressing the LNFPI biosynthetic pathway. We generated two constructs for the constitutive production of 2’FL and LNFPI in transgenic *N. benthamiana* **(Fig. 3A)**. HMO10 contains genes for the four enzymes required to produce lactose, 2’FL, LNTII, LNT, and LNFPI connected via 2A peptides^34^ to allow multiple coding sequences to be driven by a single constitutive promoter. To explore the effects of overexpressing portions of the GDP-fucose pathway, we also generated stable lines expressing HMO11, which contains a GDP-D-mannose-4,6-dehydratase (*Gmd*^24^) from the GDP-fucose pathway. *Gmd* transiently expressed alongside the neutral HMO pathway altered the HMO profile of plants in a similar way to expression of the full GDP-fucose pathway **(Supplementary Fig. 4)**. Transgenic T0 HMO-producing stable lines were assessed for fucosylated HMO yield. The majority of transgenic plants showed no drastic phenotypes compared to wildtype plants **(Fig. 3B)**. LNFPI and 2’FL were detected in leaves of transgenic *N. benthamiana* expressing both HMO10 and HMO11. Highest producing LNFPI lines accumulated an average concentration of 6.88 µg/g dry weight (dwt) **(Fig. 3C)**. Leaves from HMO11 #5 produced the highest concentration of 2’FL, reaching an average concentration of 130.35 µg/g dwt **(Fig. 3D)**. The low abundance of LNFPI in stable lines compared to transient expression could indicate that the ß-1,3-*N*-acetylglucosaminyltransferase and ß-1,3-galactosyltransferase suffer from altered activity due to the presence of 2A peptides or lower expression. Together, these results display the ability to produce and optimize a diversity of HMOs from photosynthetically-fixed CO_2_, laying the foundation for future efforts to create high HMO yielding transgenic plants for commercial HMO production.

**Figure 3.**
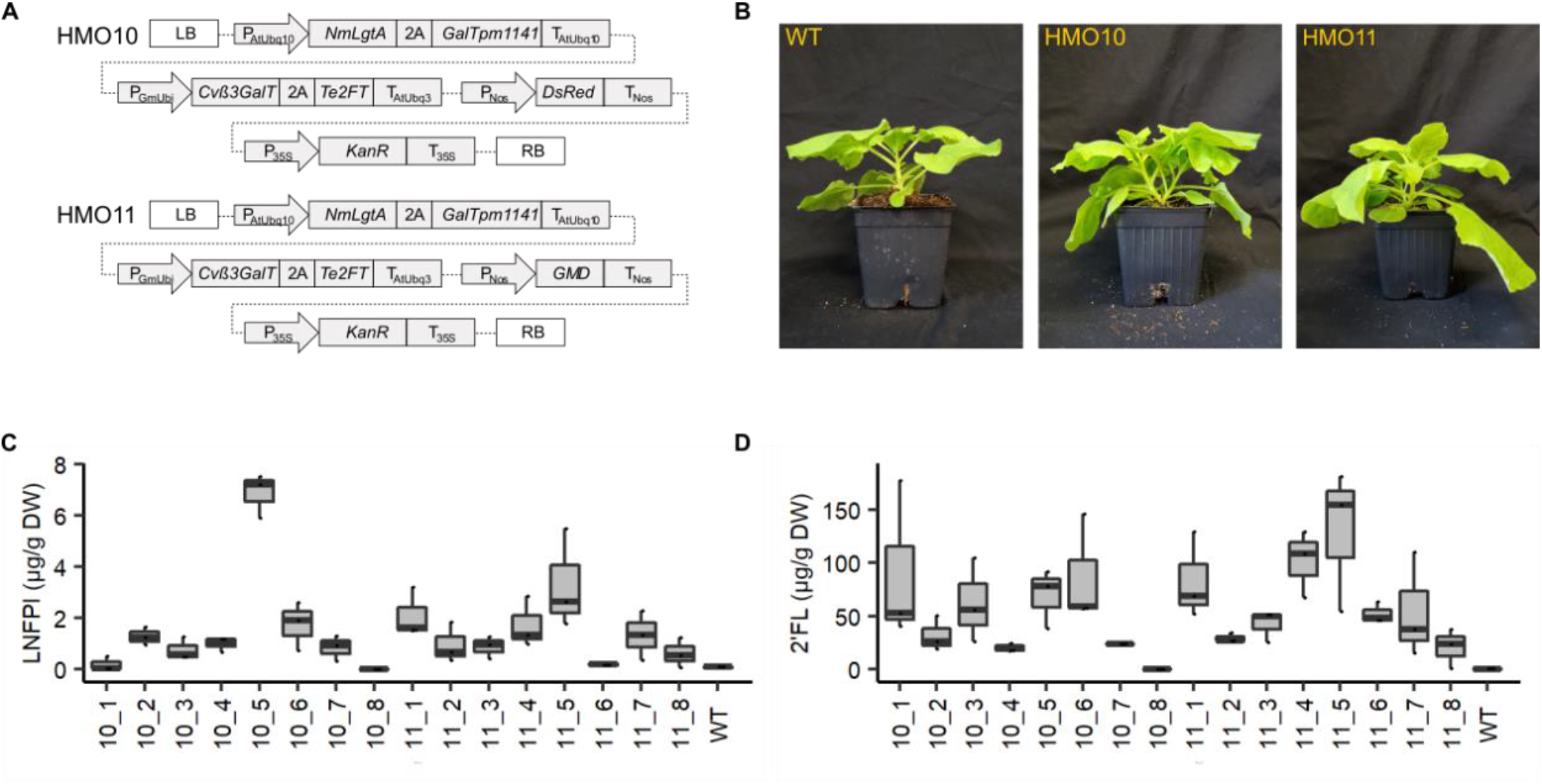
Production of human milk oligosaccharides in stably transformed plants. A) Constructs used in creation of stable lines containing biosynthetic enzymes for the production of LNFPI. B) Photos of four-week old transgenic *N. benthamiana*. C) Concentration of LNFPI produced in leaves of each stable line. D) Concentration of 2’FL produced in leaves of each stable line. For quantification, three leaves from each plant were analyzed separately. Quantification of LNFPI and 2’FL obtained with an Thermo Fisher Scientific Q-Exactive Mass spectrometer.

### Optimized purification and functional characterization of HMOs from plants

Mixtures of prebiotic sugars can have varying effects on the enrichment of beneficial gut microbes^35^. Therefore, we sought to assess the bifidogenic activity of extracts from HMO producing plants; however, crude plant extracts can contain chemicals that interfere with bacterial growth assays, such as simple sugars (glucose, fructose, sucrose) and anti-microbial phenolic compounds. Therefore, we developed a method to extract and purify HMOs from *N. benthamiana* transiently expressing the biosynthetic pathways for LNFPI **(Fig. 2A)** and GDP-fucose **(Fig. 2B)** using a novel extraction and purification process. Briefly, we performed a water extraction, yeast fermentation to remove simple sugars, and a two-step resin adsorption with Polyvinylpolypyrrolidone (PVPP) and C18 SPE. This resulted in an HMO-rich extract that contained negligible amounts of simple sugars and phenolic compounds **(Supplementary Table 3)**. The HMO extract contained target fucosylated HMOs **(Supplementary Fig. 5)**, including LNFPI, 2’FL, and LNDFHI **(Supplementary Table 4)**. The extract also contained a variety of additional oligosaccharides without assigned structures, which were composed of combinations of hexose, deoxyhexose, and HexNAc sugars **(Supplementary Fig. 5)**. These represent potentially new-to-nature oligosaccharide structures that could provide novel health benefits. Overall, these results display the ability to isolate HMOs from plants, improving their promise as an HMO production platform.

To assess the bifidogenic activity of plant-produced HMOs, we conducted growth assays to compare the effects of plant-produced HMOs to HMOs derived from human milk. We chose to assess the effects of plant-derived HMOs on *Bifidobacterium longum* subsp. *infantis* ATCC 15697 (BLI 15697) as it is a known HMO consumer^36^. We also included *Bifidobacterium animalis* subsp. *lactis* ATCC 27536 (BLAC 27536) as a negative control that does not consume HMOs^37^, but will grow on simple sugars that could be present in plant extracts or human milk. BLI 15697 grown in media containing plant-derived HMOs displayed increases in OD_600nm_ similar to that of BLI 15697 in HMO isolated from human milk, demonstrating that plant-produced HMOs possess the same selective bifidogenic activity as HMOs isolated from human milk **(Fig. 4B)**. As expected, BLAC 27536 displayed no growth either plant-derived HMOs or HMO isolated from human milk, indicating that both extracts contained a minimal amount of simple sugars present **(Fig. 4B)**. Together, these results demonstrate the ability of purified, plant-produced HMOs to mimic the bifidogenic activity of HMOs produced in humans.

**Figure 4.**
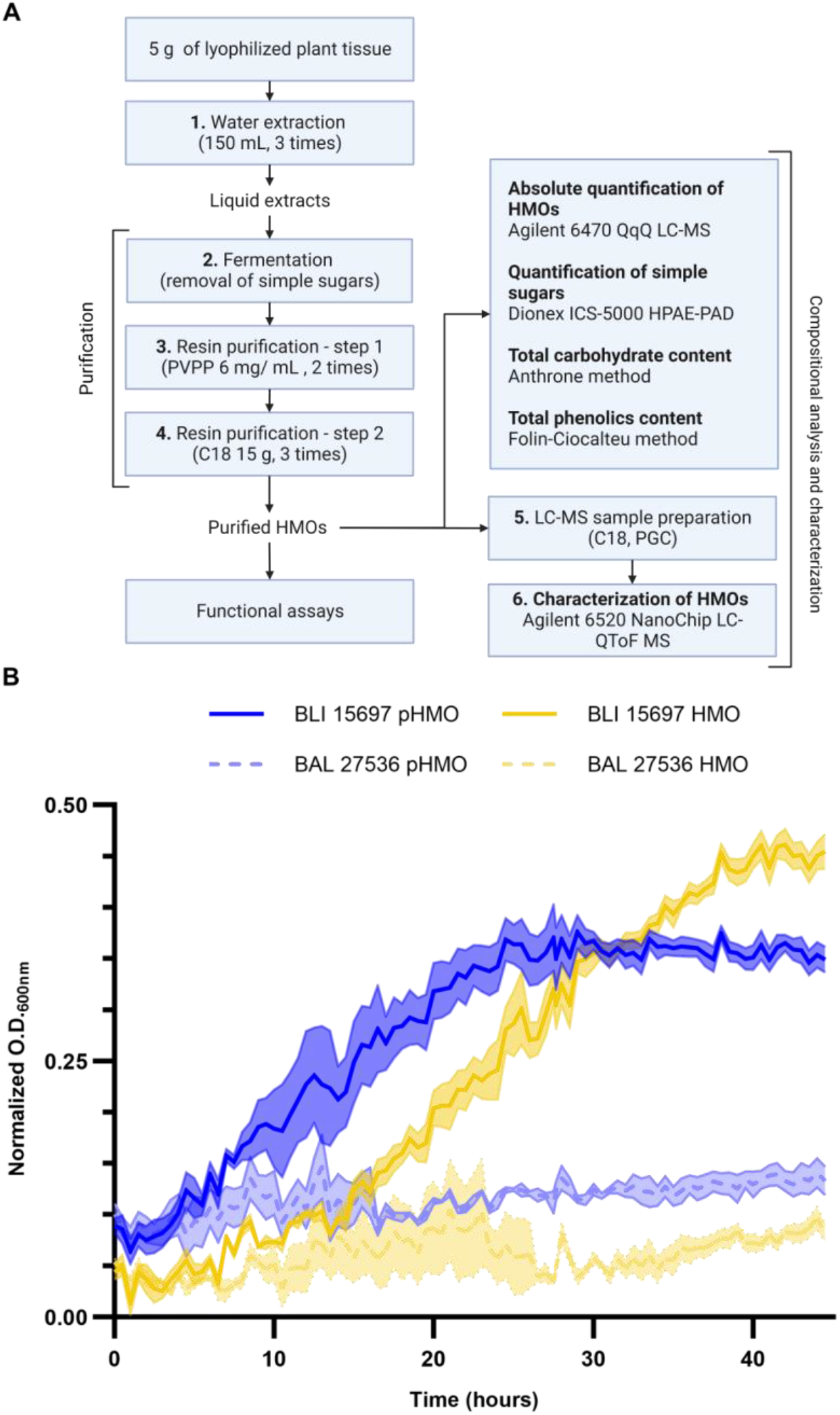
Novel purification protocol developed for functional analysis of plant-produced HMOs. A) Workflow of extraction, purification, and characterization of HMOs from *N. benthamiana* leaves. B) Growth curves of HMO consuming (BLI 15697) and control (BAL 27536) strains in media supplemented with HMO isolated from breast milk (HMO) or HMOs isolated from plants (pHMO).

### Plant-based production of HMOs are cost competitive with current microbial processes

Plant-based HMO production can be commercially viable if it demonstrates cost-competitive or cost-advantage with current state-of-the-art production routes. To assess the economic viability of HMO production in a commercially relevant crop, we developed process models and compared the cost of HMO production in plants and microbes. To do this, we performed technoeconomic analysis (TEA) of the theoretical production of LNFPI. In the plant system, we adopted typical cellulosic biorefinery design utilizing biomass from sorghum to coproduce HMOs along with biofuel because co-producing value-added bioproducts in biorefineries is a promising approach to maximize the utilization all of biomass and hence improve the economics of biorefineries^38,39^. We assume that biomass sorghum can accumulate 0.31% dry weight of LNFPI in the entire biomass, as this was our highest yield following purification of LNFPI **(Supplementary Table 4)**. We also developed process models and conducted TEA for the HMOs in *E. coli* using the established processes and the highest reported yields of LNFPI^40^ from peer-reviewed papers at the time of conducting this TEA. The comparison between plant and microbial systems to produce the same product aims to provide in-depth understanding of the cost benefits of the systems.

TEA results indicate that producing LNFPI from the plant system is economically favorable compared to the microbial system **(Fig. 5)**. In the plant system, the minimum selling prices of LNFPI are $5.2/kg when selling ethanol at the cellulosic ethanol selling price and $18.9/kg when ethanol is sold at the target fuel price, respectively **(Fig. 5A)**. However, microbial-based LNFPI results in a minimum selling price (MSP) of $85.5/kg with glucose being the largest cost contributor **(Fig. 5B)**. The high cost obtained in the microbial system is largely due to its extremely low highest-reported yields (0.02%) and recovery rate after bioconversion (62%). These results indicate that when microbial hosts are unable to produce HMOs at a comparable yield, plants may be a cost-effective bioplatform for high-value products. In addition to relative cost advantages of plant systems, using biomass as the feedstock to co-produce high-value compounds and biofuel offers considerable environmental benefits because biomass can remove CO_2_ from the atmosphere^41^. Although current HMO yields in stable lines of the model plant, *N. benthamiana* **(Fig. 3)** are below the yields in transiently expressing tissue, high yields could be achieved by optimizing the HMO constructs used for the production of stable lines, finding an optimal crop species for HMO production, and optimizing growing conditions.

**Figure 5.**
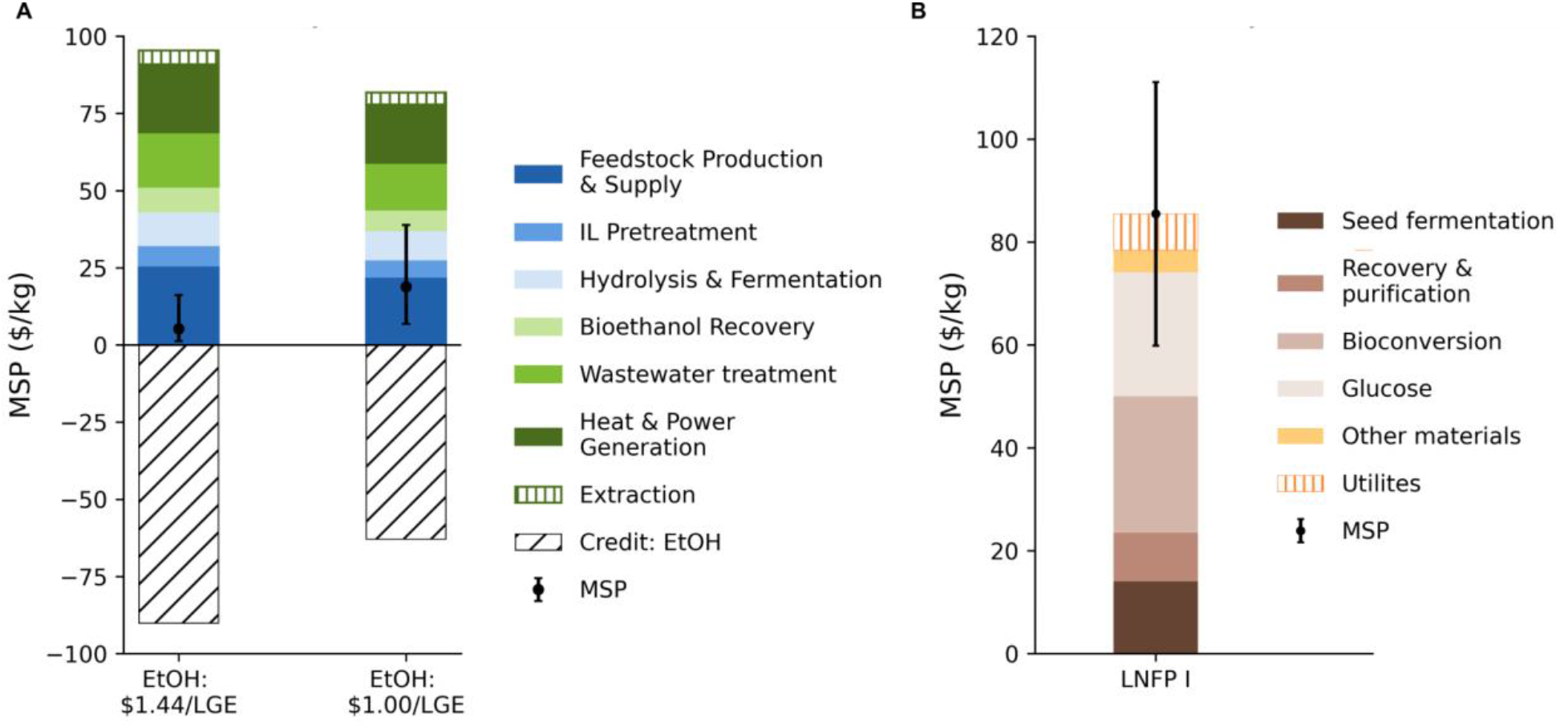
Plant-based production improves the economics of LNFPI production. A) Estimated minimum selling price (MSP) of LNFPI produced using biomass sorghum as a model production platform. B) Estimated MSP of microbially produced LNFPI based on inputs, yield, and process described in the highest microbial production of LNFPI^40^.

## Discussion

Human milk oligosaccharides are major contributors to the bioactive properties of human human milk^1,2^. While the unique bioactivities of a small number of HMOs have been investigated^3,7–10^, over 100 HMOs remain to be studied due to lack of access to material, representing a wealth of potential bioactive molecules. Here, we report the production of all general three classes of human milk oligosaccharides *in planta*. Notably, expression of HMO biosynthetic pathways *in planta* created a variety of complex HMOs including oligosaccharides that currently cannot be produced in microbial platforms, which indicates plants could serve as a platform for the production of a range of HMOs that are not currently producible in microbial hosts. Furthermore, we optimized the production of both specific HMOs and HMO classes by overexpressing nucleotide sugar biosynthetic pathways. The diversity of HMOs produced from inserting a relatively small number of genes displays the ability of plants to generate complex sugars. Transient expression serves as a viable platform for testing HMO biosynthetic genes and small scale production of HMOs for functional validation. Novel, lab-scale purification methods enabled the isolation of oligosaccharides, including HMOs, from plant tissue, displaying the potential of plants as an industrial source of HMOs. HMOs purified from plants provided selective bifidogenic activity, indicating they would serve as a potent prebiotic *in vivo.* Future pathway engineering could yield biosynthetic pathways for the production of all HMOs present in human milk, including complex, branched HMOs. This would enable the study and potential consumption of bioactive HMOs that are currently unavailable.

Despite the field of plant synthetic biology being in its nascent stages, plants have the potential to produce compounds of interest at lower costs than microbial platforms due their intrinsic and unique metabolic capabilities^38^. Additionally, plants are capable of using atmospheric carbon dioxide during their growth cycle to produce target compounds and biofuels, improving the sustainability of target compound production. We demonstrate the ability of stably transformed plants to produce HMOs that are commercially available (2’FL) and currently unavailable (LNFPI). Our technoeconomic analysis results indicate that HMO production in commercially relevant crops could be a more cost-effective platform than microbial production for both simple and complex HMOs.

As the demand for HMOs increases due to the growing infant formula and adult prebiotic markets, plants will serve as a novel platform for the production of diverse HMOs at agricultural scales. Furthermore, the diversity of plant-produced HMOs will provide researchers with access to HMOs that were previously inaccessible. Since the structure of an HMO determines its bioactivity, this could lead to the discovery of HMOs that treat various gastrointestinal illnesses. Additionally, production of HMOs in plants could permit direct consumption as food by directly ingesting the plant or products made from the plant. Such a product may serve as a consumable source of prebiotic HMOs for humans or be added to forage crops for animal consumption. Overall, the production of HMOs *in planta* provides the opportunity to simultaneously improve the scale of HMO production and expand the diversity of HMOs available to improve the gastrointestinal health of infants and adults.

## Methods

### Plasmid construction and transient expression

For transient expression, candidate glycosyltransferases were PCR amplified and cloned into the binary vector, PMS057^42^, using Golden Gate assembly^43^, Gibson assembly^44^, or restriction-ligation. XL1-blue *Escherichia coli* cells were transformed with the assembled plasmids via heat shock^45^. Transformed cells were selected by plating cells on Lysogeny broth (LB) agar plates containing 50 µg/mL kanamycin. Plasmid assembly was confirmed via miniprep and Sanger sequencing (Azenta). *Agrobacterium tumefaciens* str. GV3101 was transformed using sequence-verified plasmids by electroporation^46^. Transformed colonies were selected using LB agar plates containing 50 µg/mL kanamycin, 50 µg/mL rifampicin, and 10 µg/mL gentamicin. *A. tumefaciens* str. GV3101 harboring individual candidate glycosyltransferases were grown in LB overnight to an OD_600_ (VWR, V-1200) of 0.8 to 1.2. The cultures were centrifuged at 4000 x g for 10 min, and the supernatant was decanted. Cell pellets were resuspended in infiltration media (10 mM MES, 10 mM MgCl_2_, 500 µM acetosyringone, pH 5.6) and incubated at room temperature for one hour with gentle rocking (Thermolyne, VariMix). *A. tumefaciens* strains harboring each glycosyltransferase were mixed in equal amounts alongside a strain harboring the p19 silencing suppressor^47^ to reach a final OD_600_ of 0.5. *A. tumefaciens* mixtures were injected into the abaxial side of a leaf on a four-week-old *Nicotiana benthamiana* using a needleless syringe. Each experiment was performed with three biological replicates.

For the production of stable lines, HMO10 and HMO11 constructs were generated through a multi-part Golden Gate assembly containing subcloned transcriptional units. Assembled plasmids were transformed and sequence verified as described above. *N. benthamiana* was transformed using *A. tumefaciens* str. EHA105 harboring HMO10 or HMO11 by the UC Davis Plant Transformation Facility.

### HMO extraction for identification of HMOs from individual leaves

*N. benthamiana* leaves transiently expressing HMO biosynthetic enzymes were harvested five days post infiltration. Three leaves of *N. benthamiana* stable lines transformed with HMO10 and HMO11 were harvested at four weeks old. Following harvest, vasculature was removed, and leaves were frozen in liquid nitrogen and lyophilized (Labconco, Freezone 4.5) for two days. Lyophilized leaves were homogenized via a bead mill (Retsch, MM400) at 20 Hz for 10 minutes. Oligosaccharides were extracted from 20 mg of lyophilized leaf tissue by ethanol precipitation. To each sample, 1 mL of 80% ethanol was added before homogenization on a bead mill at 10 Hz for 1 min. Samples were then precipitated overnight at -20 °C and centrifuged at 10,000 rpm for 15 min. The supernatant was transferred to a 2 mL screw-cap tube. The pellet was washed twice by adding 500 μL of 80% ethanol, homogenizing via bead mill for 1 min, and centrifuging at 10,000 rpm for 15 min. The supernatant and washes were combined and dried in a vacuum centrifuge (Genevac EZ-2, SP Scientific). Dried supernatants were reconstituted in 200 μL of water and subjected to both C18 and porous graphitized carbon (PGC) solid phase extractions (SPE) (Thermo) in 96-well plate format. C18 cartridges containing 25 mg of stationary phase were first conditioned by two additions of 250 μL of acetonitrile (ACN) followed by four additions of 250 μL water. Samples were then loaded and eluted with two volumes of 200 μL water. PGC cartridges containing 40 mg of stationary phase were conditioned by addition of 400 μL water, 400 μL 80% (v/v) ACN and water, followed by two volumes of 400 μL water. The sample eluate from C18 SPE was then loaded, washed thrice with 500 μL water, and eluted using two volumes of 200 μL 40% (v/v) ACN and water. The purified extracts were dried in a vacuum centrifuge and reconstituted in 100 μL of water before injecting 5 μL for liquid chromatography mass spectrometry (LC-MS) analysis.

### LC-MS analysis of HMOs from individual leaves

For initial screening, chromatographic separation was carried out using a Thermo Scientific Vanquish UHPLC system equipped with a Waters BEH C18 Amide column (HILIC) (1.7 µm, 100 mm x 2.1 mm). A 10 min binary gradient was employed based on Xu et al. (2017)^48^: 0.0 – 4.0 min: 25-35% A; 4.0 – 8.50 min, 35-65% A; 8.50 – 8.70 min: 25% A. Mobile phase A consisted of 3% ACN (v/v) in water with 0.1% formic acid, and mobile phase B consisted of 95% ACN (v/v) in water with 0.1% formic acid.

For identification of HMOs produced, LC-MS analysis was performed using a Thermo Scientific Vanquish 3000 UPLC system connected to Thermo Scientific qExactive mass spectrometer. Chromatographic separation was carried out using a Hypercarb porous graphitic carbon (PGC) column (5 µm, 150 mm x 1 mm, Thermo Scientific). A 40-minute binary gradient using 3% ACN in water containing 0.1% formic acid (Solvent A) and 90% (v/v) ACN in water containing 0.1% formic acid was performed as follows: 100% A, 0 - 2.5 min; 100 - 84% A, 2.5 - 15 min; 84 - 42% A, 15 - 20 min; 42 - 0% A, 20 - 22 min; 0% A, 22 - 28 min; 0 - 100% A, 28 – 30 min; 100% A, 30 - 40 min.

For identification of HMOs, the Q-Exactive mass spectrometer equipped with an electrospray ionization source was operated in positive ionization mode with the following parameters: scan range = 133.4 - 2000 *m/z*, spray voltage = 2.5 |kV|, capillary temperature = 320°C, aux gas heater temperature = 325°C, sheath gas flow rate = 25, aux gas flow rate = 8, sweep gas flow rate = 3. MS/MS analysis was performed using stepped collision energies of 20, 30, 40 [eV]. MsDIAL was used for data analysis^49^.

For quantification of LNFPI and HMO profiling, mass spectral analysis was carried out on an Agilent 6530 Accurate-Mass Q-TOF MS operated in positive mode using data dependent acquisition. The gas temperatures were held at 150 °C. The fragmentor, skimmer, octopole, and capillary were operated at 70 V, 55 V, 750 V, and 1800 V, respectively. The collision energy was based on the empirically derived linear formula (1.8 × (m/z/100) - 3.6). The reference mass used for calibration was 922.009798 m/z. The Agilent MassHunter Qualitative software was used for data analysis. Oligosaccharides were identified using an in-house library, their MS/MS spectra, and comparison to either authenticated standards or a pool of human milk oligosaccharides of known composition.

### Extraction and purification of HMOs from pooled leaves

Five grams of lyophilized and ground *N. benthamiana* leaves transiently expressing the LNFPI and GDP-fucose biosynthetic pathways was mixed with 150 mL of water and agitated for 15 min at room temperature in a stirring plate. The mixture was centrifuged at 4000 x g for 5 min, and the supernatant was separated. The extraction was repeated two more times, combining the supernatant each time. The final supernatant was filtered using a 0.22 µm Millipore Steritop vacuum filter. The extraction process was carried out in duplicate to ensure reproducibility.

Yeast fermentation was carried out to eliminate simple sugars (glucose, sucrose, and fructose) from the extracts^50^. Briefly, autoclaved extracts were inoculated with 0.4 g/L of commercial active dry yeast *Saccharomyces cerevisiae* (UCD 522 Montrachet, Lallemand Inc., Montreal, Canada) at 30 °C, 150 RPM for 24 h. After 24 hours, the samples were centrifuged at 4000 x g for 5 minutes and filtered using a 0.22 µm Millipore Steritop vacuum filter to remove the yeast. Samples were concentrated using a vacuum concentrator (Genevac™ miVac Centrifugal Concentrator (Ipswich, UK) at room temperature and frozen until their purification.

PVPP (Sigma-Aldrich, St. Louis, MO) was used to bind phenolic compounds within the sample following a previous protocol^51^. Briefly, 3 g of PVPP was conditioned by mixing it with 100 mL of 12 M HCl at 100 °C for 30 min with constant stirring in a stirring plate. After cooling off, the slurry was centrifuged at 4000 x g for 5 min and filtered using a 0.22 µm Millipore Steritop vacuum filter. Subsequently, the PVPP was washed with nanopure water until the flow-through reached a pH of 7. Activated PVPP was mixed with water to a final concentration of 20 mg PVPP/mL.

PVPP suspension was added to the extracts at a concentration of 6 mg PVPP/mL and agitated at room temperature for 15 min on a stirring plate. After the time had elapsed, the sample containing the plant extracts and the PVPP was centrifuged at 4000 x g for 5 min and filtered using a 0.22 µm Millipore Steritop vacuum filter to separate the PVPP containing the bound phenolics. To eliminate residual phenolics from the extracts, additional PVPP was added to the supernatant (6 mg PVPP/mL), and the process was repeated. The filtrate containing the oligosaccharides was concentrated and frozen until further purification.

Two SPE columns of 60 mL were packed with 15 g of Bondesil – C18, 40 µm suspended in 20 mL of ACN. After the ACN was drained, a frit was added to the C18. Before loading the samples, the columns were conditioned with 3 volumes of ACN and 3 volumes of nanopore water. PVPP-treated extracts were loaded onto the conditioned C18 columns, and the oligosaccharides were washed with 250 mL nanopure water divided into four washes. To ensure the complete removal of interfering compounds, C18 SPE was repeated 2 more times. The purified HMO fractions were dried in a vacuum concentrator (Genevac™ miVac Centrifugal Concentrator (Ipswich, UK) at room temperature and frozen.

### Compositional analysis of plant material

Total carbohydrate content was assessed by the Anthrone method with modifications^52^. In a 96-well microplate, 40 μL of purified and diluted extracts were combined with 100 μL of Anthrone reagent (2 mg/mL (w/v) in cold 98% sulfuric acid) and mixed through pipette tip aspiration. The microplate was incubated for 3 min at 92 °C in a water bath followed by 5 min at a room temperature water bath and then for 15 min in a 45 °C Thermolyne™ Benchtop muffle furnace (Thermo Fisher Scientific, Waltham, MA, USA). The plate was cooled for 3 min at room temperature before measuring the absorbance with a SpectroMax M5 UV/Vis spectrophotometer (Molecular Devices, San Jose, CA, USA) at 630 nm. Total carbohydrate quantification calculations were based on a glucose standard curve. Each plant extract was prepared in duplicate, and each sample was further analyzed in duplicate.

Total phenolic content of the extracts was determined according to the Folin-Ciocalteu spectrophotometric method as described by Singleton et al.^53^

Simple sugars (glucose, sucrose, and fructose) were quantified by high-performance anion exchange chromatography with pulsed amperometric detection on a ThermoFischer Dionex ICS-5000+HPAE-PAD system based on a method described by Huang et al. with modifications^54^. Diluted extracts (1:100, v/v, or 1:1000 in nanopure water) were filtered through a 0.2 mm syringe filter (Agilent Captiva Econo Filter, PES, 13mm, 0.2 μm) into 2 mL vials with septa. The samples (25 μL) were injected into a CarboPac PA200 guard column (3 x 50 mm) and a CarboPac PA200 analytical column (3 x 250 mm), and chromatographic separation was carried out with a 12 -min gradient elution (from 0.6 to 25% B in 12 min), 0.5 mL min^-1^ flow rate. The solvent system consisted of A: 100% water and B: 200 mM sodium hydroxide (NaOH). Calibration curves (correlation coefficient ≥ 0.999) were prepared using glucose, sucrose, and fructose standards.

### Quantification of HMOs by QqQ LC-MS

Detection and quantitation of HMOs were performed using an Agilent 6470 Triple Quadrupole Liquid Chromatography Mass Spectrometry System (QqQ LC-MS) equipped with an Advance Bio Glycan Map column (2.1 × 150 mm, 2.7 μm, Agilent). The mobile phase consisted of 10 mM ammonium acetate in 3 % ACN, 97 % water (v/v, pH 4.5; A) and 10 mM ammonium acetate in 95 % ACN, 5 % water (v/v, pH 4.5; B). The chromatographic separation was carried out at 35 °C with gradient elution at a flow rate of 0.3 mL/min. The MS analysis was conducted in positive ion mode with source parameters as follows: the gas temperature was 150 °C at a flow rate of 10 L/min; the nebulizer was 45 psi; the sheath gas temperature was 250 °C at a flow rate of 7 L/min; capillary voltage was 2200 V. Please see Supplementary Table 2 for gradient and multiple reaction monitoring (MRM) transitions.

### Characterization of HMOs by LC-QToF-MS

Oligosaccharides were purified by a two-step solid-phase extraction (SPE) using C18 (HyperSep C18-96, 50 mg bed weight; Thermo Fisher Scientific) and porous graphitic carbon (PGC) (HyperSep Hypercarb-96, 25 mg bed weight; Thermo Fisher Scientific)^55^. The samples were filtered (Captiva Premium Syringe Filter, polyethersulfone (PES) membrane, 4 mm diameter, 0.2 µm pore size, LC/MS certified) into 200 µl vials.

Individual oligosaccharide compositions were analyzed with an Agilent 6520 NanoChip LC-QToF mass spectrometer (Santa Clara, CA, USA). Oligosaccharides separation was achieved with a microfluidic high-performance liquid chromatography (HPLC) PGC chip containing an enrichment (4 mm, 40 nL) and an analytical (75 μL x 43 mm) column as well as a nanoelectrospray tip, using a binary solvent gradient of solvent A (5 mM ammonium acetate in 3% ACN, 97% water (v/v)) and solvent B (5 mM ammonium acetate in 90% ACN, 10% water (v/v)) based on a previously optimized method^54^. The gradient was 0–16% B at 0–20 min, 16–44% B at 20–30 min, 44–100% B at 30–35 min, 100% B at 35–45 min, and 100–0% B from 45 to 45.1 min, followed by a 15 min re-equilibration of 100% A^56^. The mass spectrometer was operated in positive ionization mode with a range of m/z 320-2500 and an electrospray capillary voltage of 1800-1900 V. Reference masses of m/z 922.009 and 1221.991 provided continuous internal calibration. All samples were analyzed using tandem mass spectrometry (MS/MS) with tandem fragmented peaks selected by the automated precursor selection of the six ions with highest signal intensity with a medium isolation width. The QToF MS had a ramped collision energy slope of 0.02 based on m/z values with an offset of -3.5 V. The acquisition rate of 1 spectrum/s was used for both MS and MS/MS. Each spectrum was manually examined, and molecular masses were confirmed with Agilent MassHunter Qualitative Analysis B.07.00 software using the molecular feature extraction and a maximum tolerance of 20 ppm.

### Bacterial strains and growth conditions

*Bifidobacterium longum* subsp. *infantis* ATCC 15697 and *Bifidobacterium animalis* subsp. *lactis* ATCC 27536 were cultured at 37°C in a Coy vinyl anaerobic bubble with an atmosphere of 2.5% H2, ∼5% CO_2_, and balance N_2_. Routine culturing was done with Difco MRS + 0.05% L-cysteine HCl (MRSC), and carbohydrate-specific culturing was done with modified MRS (mMRSC), which was prepared per liter as follows: 10 g Bacto proteose peptone #3, 10 g Bacto casitone, 5 g Bacto yeast extract, 2 g triammonium citrate, 5 g sodium acetate trihydrate, 200 mg magnesium sulfate hexahydrate, 34 mg manganese sulfate monohydrate, 0.5 g L-cysteine HCl, and 1.063 g Tween-80. Normally, 2 g anhydrous dipotassium phosphate would also be added, but it was precipitated by the plant HMO preparation.

### Growth curves

1 mL 1 mMRSC + dipotassium phosphate + 1% lactose monohydrate was inoculated with one colony and incubated for 24 hours. The growth curve inoculum was cultured by diluting the 24-hour culture 1:100 in mMRSC + dipotassium phosphate + 1% lactose, then incubating for 12-15 hours. The inoculum was prepared by washing the cells twice with one volume of 1x PBS. 160 ul growth curve cultures were contained in flat-bottomed, optically clear, 96-well, lidded plates, and they had a final inoculum and sugar concentration of 1% in mMRSC. Lactose was the growth control substrate, water was the no-growth control substrate, and pooled HMO^57^ was the HMO-growth control substrate. Cultures were done in triplicate. Wells were overlaid with 40 ul sterile mineral oil and incubated in a BMG SpectroStar Nano. The plate reader was set to read the O.D.600nm of each well 30 times in a spiral pattern every half hour with medium orbital shaking before each read. All media had uninoculated controls whose O.D.600nm was subtracted from that of the inoculated medium.

### Technoeconomic analysis

In this study, we used *SuperPro Designer v12* to develop technoeconomic models of HMOs that can be produced from both plant and microbial systems. To do this, we first developed process models and then applied discount cash flow analysis of the theoretical production of LNFPI in plants and microbes. The simplified process flow diagram can be found in **Supplementary Fig. 6**. In the plant system, we adopted integrated cellulosic biorefinery design to co-produce HMO and ethanol to maximize the utilization of plant biomass. Biomass sorghum was used as the representative plant because its characteristics such as high yields and drought tolerance are ideal for biofuel production. Previous studies demonstrated that using biomass sorghum as the representative plant to accumulate value-added bioproducts and bioethanol, as the representative fuel, produced could improve the performance of an integrated biorefinery^38,39^. Since ethanol is co-produced in the biorefinery, two ethanol selling prices were considered: 1) baseline cellulosic biofuel selling price of $1.44/liter of gasoline equivalent (LGE) and 2) target fuel price of $1.00/LGE.

Briefly, biomass sorghum with 0.31% dwt HMOs accumulated in the plant biomass are harvested and transported to the biorefinery gate for pre-processing and short-term on-site storage. HMO extraction is then conducted based on our lab process; the extracted HMOs are separated, purified, and recovered as the main product, the remaining biomass from biomass sorghum is routed to ionic-liquid (IL) pretreatment for biomass deconstruction. After IL pretreatment, enzymatic hydrolysis and ethanol fermentation is conducted to produce ethanol, followed by distillation and molecular sieve to remove excess water. Wastewater from the overall process is routed to the wastewater treatment sector to produce reusable process water and biogas, which can be combusted in the onsite turbogenerator, along with other solids from biomass, to generate heat and electricity that can satisfy the facility’s need. In the microbial system, HMO was produced as the single product in the biorefinery. Unlike the plant system, pure sugar is used in the microbial system as the sole feedstock and no wastewater treatment sector and onsite combustion are designed for the microbial system.

After developing technoeconomic models in *SuperPro Designer*, we performed mass and energy balance, and then applied discounted cash flow analysis to quantify the minimum selling price of HMOs ($/kg). For both systems, we assume the biorefinery can operate 24 hours per day and 330 days per year for 30 years. The unit price of biomass sorghum is assumed at $95/bone-dry tonne and we assume in the plant system, the cellulosic biorefinery can intake 2,000 bone-dry tonnes of biomass sorghum per day. The unit price of IL is $2/kg with a range of $1/kg to $5/kg. Other parameters are kept constant as the 2011 NREL report^58^.

## Acknowledgements

CRB acknowledges support from the National Institutes of Health NIGMS T32 Training Program and the Department of Energy. CRB and PMS acknowledge support from the DOE Joint BioEnergy Institute (http://www.jbei.org) supported by the U. S. Department of Energy, Office of Science, Office of Biological and Environmental Research, through contract DE-AC02-05CH11231 between Lawrence Berkeley National Laboratory and the U.S. Department of Energy. PMS acknowledges support from grant number R00AT009573 from the National Center for Complementary and Integrative Health (NCCIH) at the National Institutes of Health. MY would like to acknowledge USDA NIFA HATCH NC02948. DAM acknowledges support of the Peter J. Shields Chair in Dairy Food Science. DB acknowledges the Hatch project CA-D-FST-2744-H. We would like to thank Dr. Xi Chen for sharing plasmids with genes encoding several glycosyltransferases.

## Supplementary Materials for

### Supplementary Figures

**Supplementary Figure 1.**
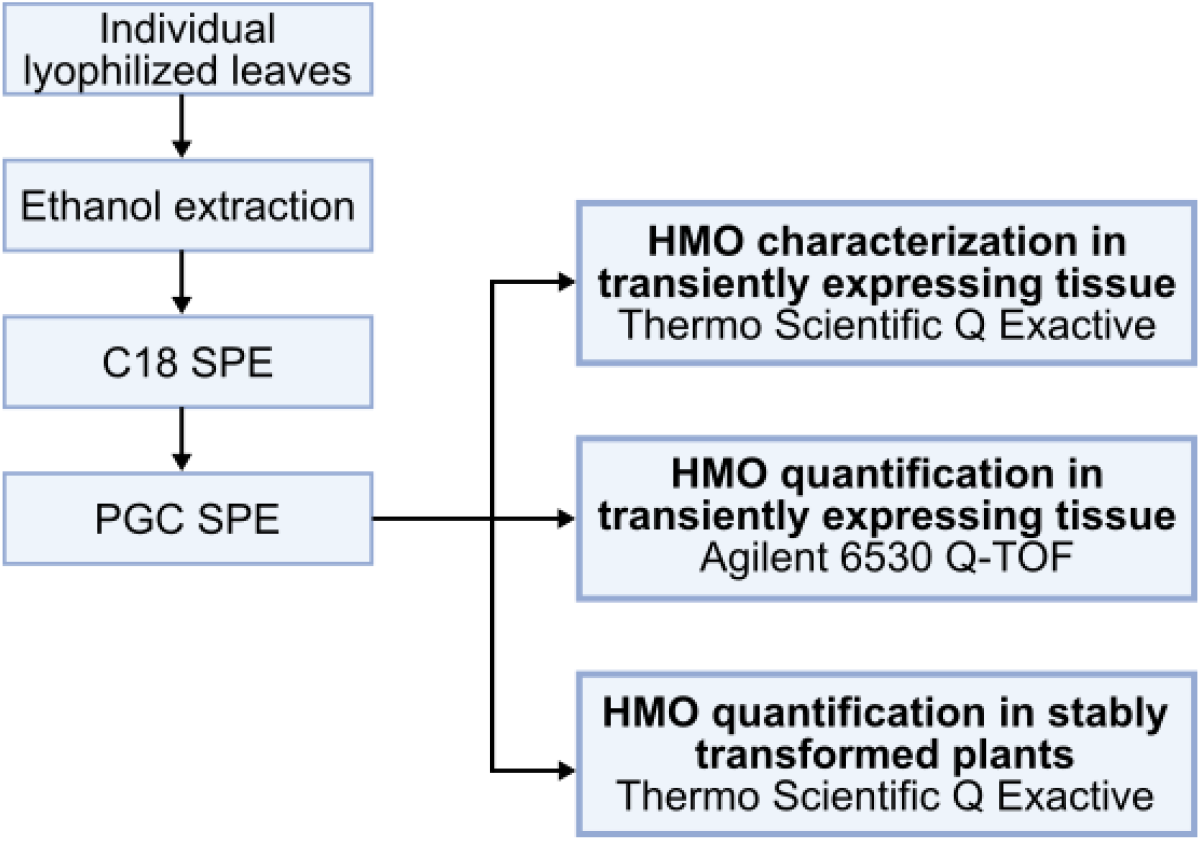
**Flow diagram displaying workflows for the characterization and quantification of HMOs from single leaves transiently expressing HMO biosynthetic pathways.**

**Supplementary Figure 2.**
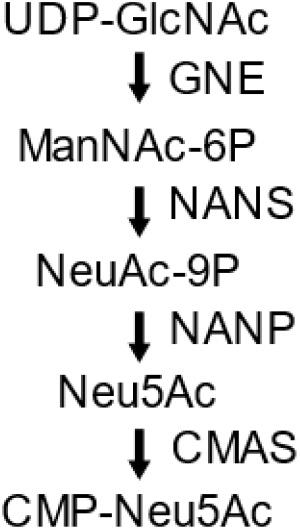
**Mammalian CMP-Neu5Ac pathway used for production of acidic HMOs.**

**Supplementary Figure 3.**
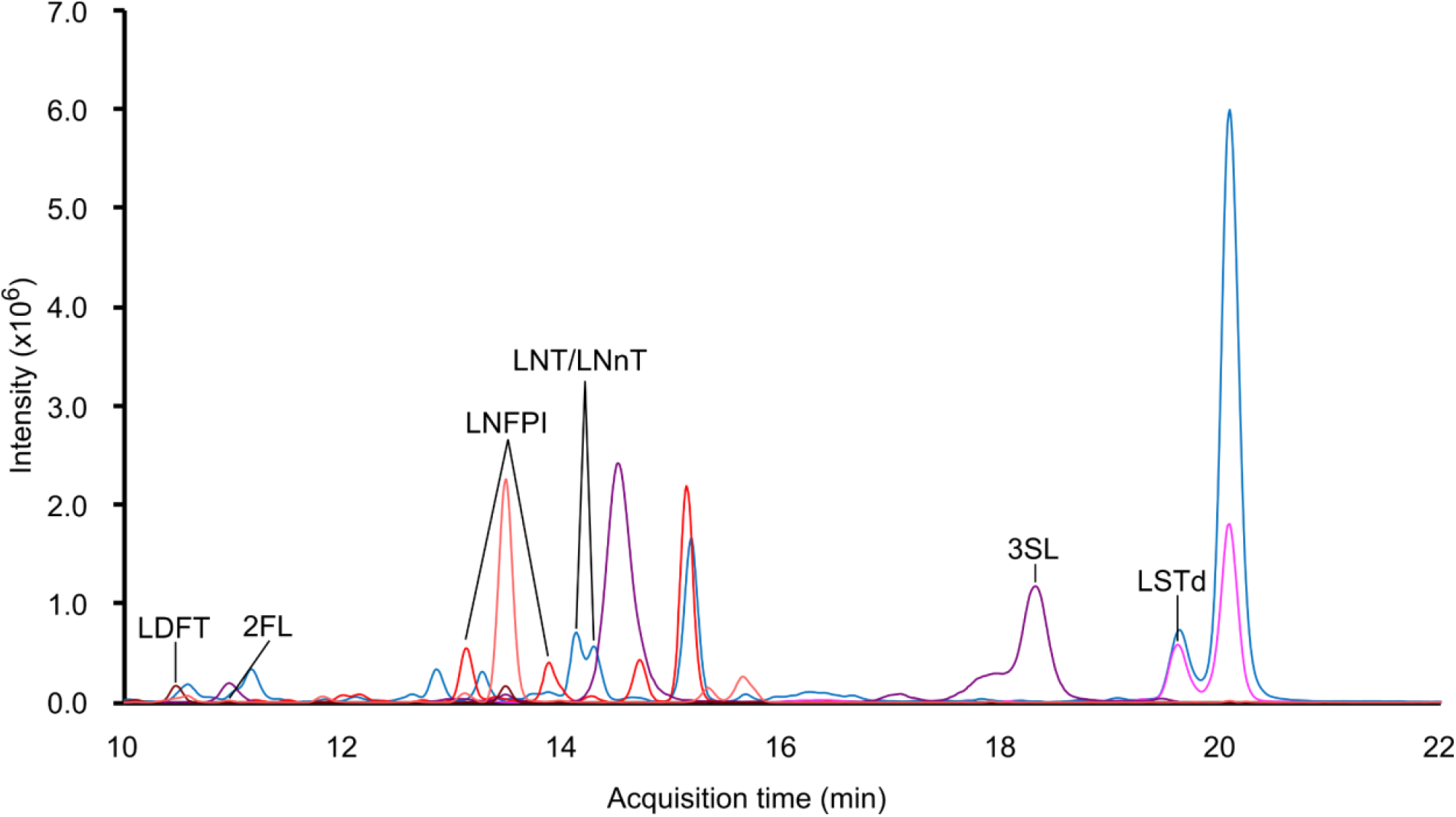
Production of all three HMO classes in a single plant leaf. Stacked extracted ion chromatogram obtained using Thermo Fisher Scientific Q-Exactive mass spectrometer showing identification of HMOs produced in a single leaf expressing the genes for production of neutral, fucosylated and sialylated HMOs (*GalTpm1141*, *NmLgtA*, *Cvß3GalT*, *Hp0826*, *Te2FT*, *PmSt3*, *St6*, *GNE*, *NANS*, *NANP* and *CMAS*). Additional peaks represent in-source fragments of larger oligosaccharides or additional isomers. Chromatographic separation was performed using a porous graphitic carbon column. All labeled peaks were identified using analytical standards.

**Supplementary Figure 4.**
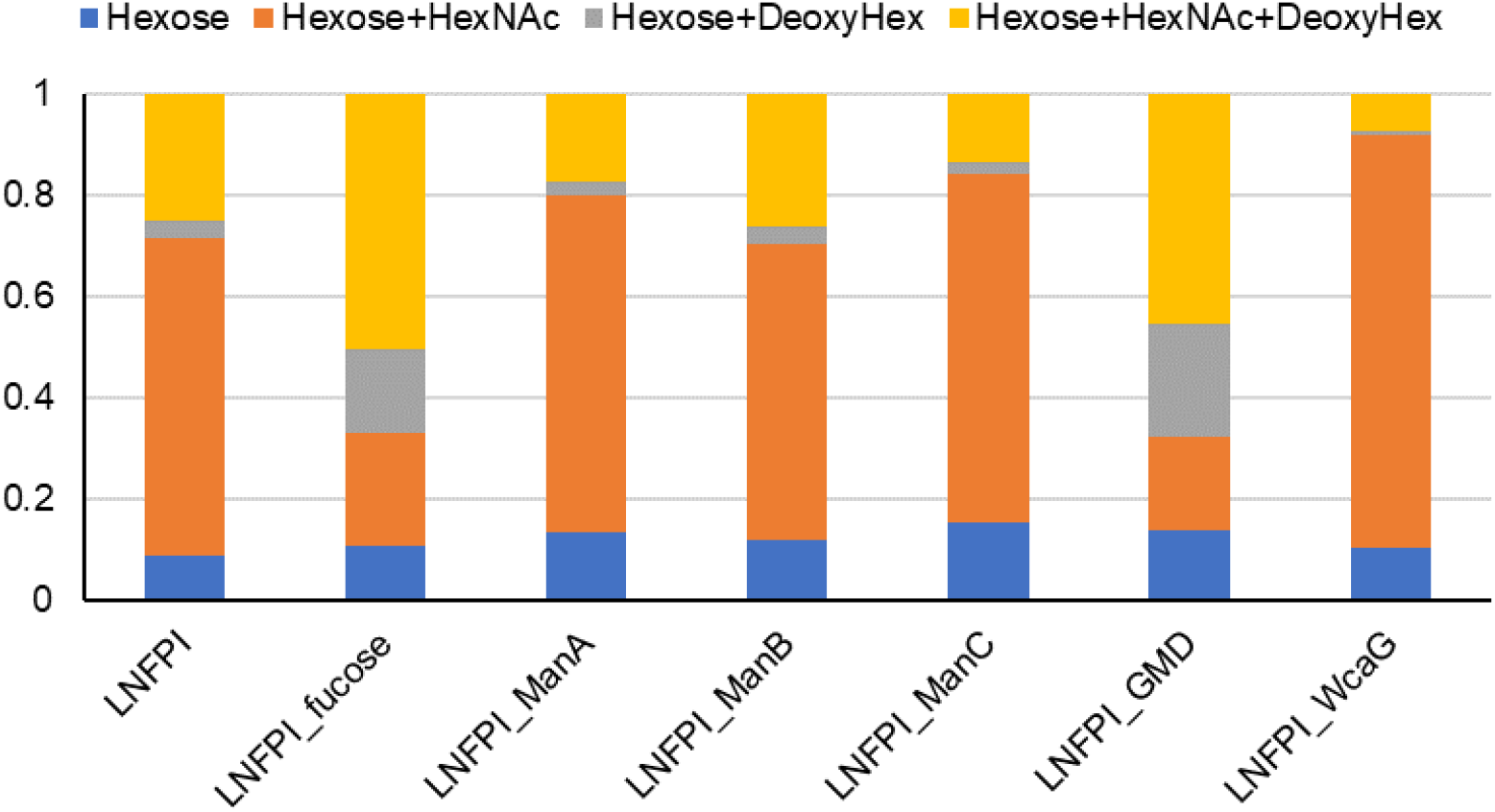
Effect of overexpression of individual enzymes from the GDP-fucose biosynthetic pathway on HMO profile produced using the LNFPI pathway. Composition based on Hexose, HexNAc, Deoxyhexose (Deoxyhex) composition determined using *m/z* and MS/MS fragmentation. Obtained with an Agilent 6530 Accurate-Mass Q-TOF MS.

**Supplementary Figure 5.**
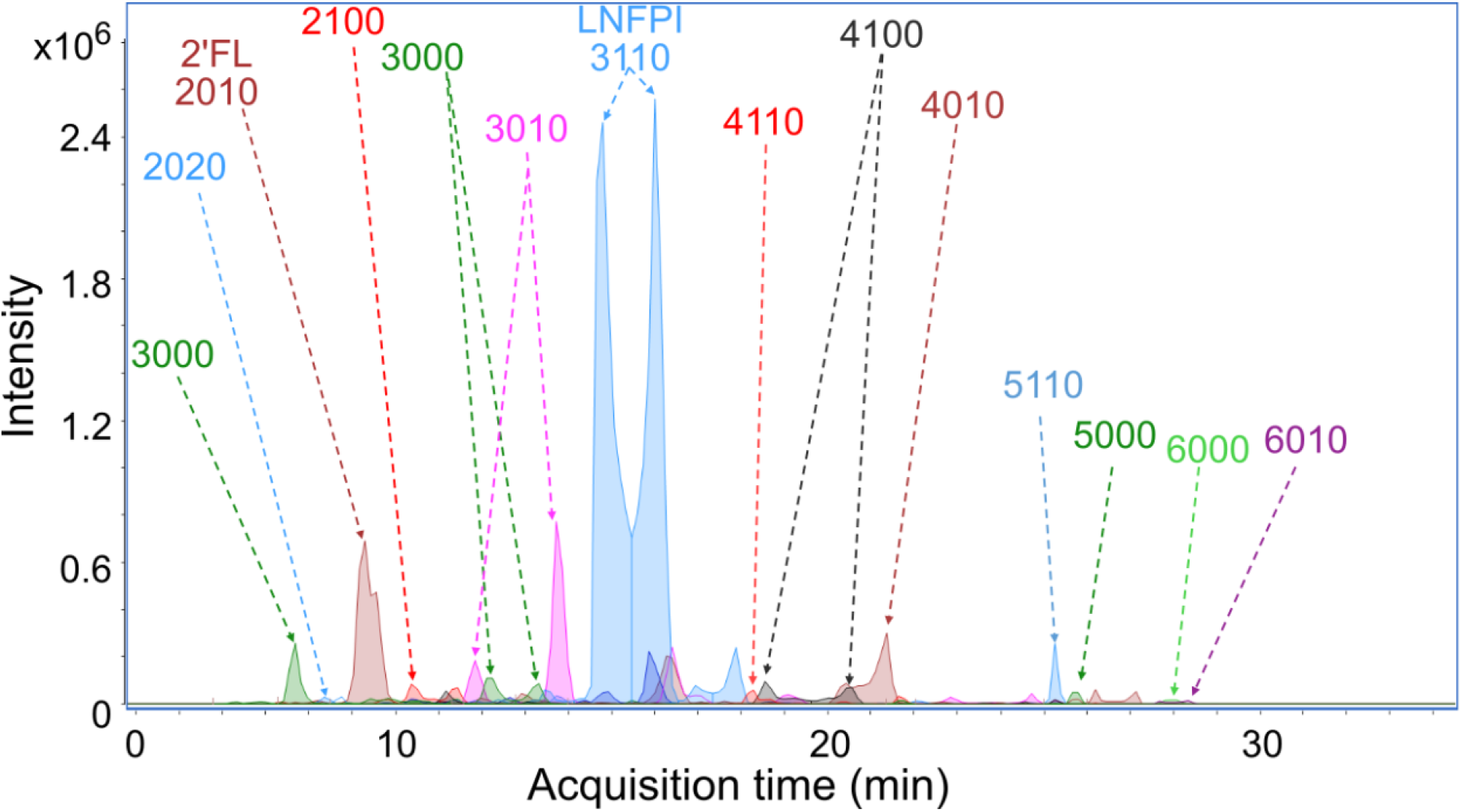
Representative Extracted Compound Chromatogram (ECC) for *N. benthamiana* oligosaccharides produced from one of the *N. benthamiana* replicates, obtained with an Agilent 6520 NanoChip LC-QToF. Oligosaccharides names are annotated based on their composition expressed as a numerical in which the first digit represents the number of hexose sugars, second digit indicates the number of hexNAc sugars, third digit indicates the number of deoxyhexose sugars, and the fourth digit represents the number of Neu5Ac.

**Supplementary Figure 6.**
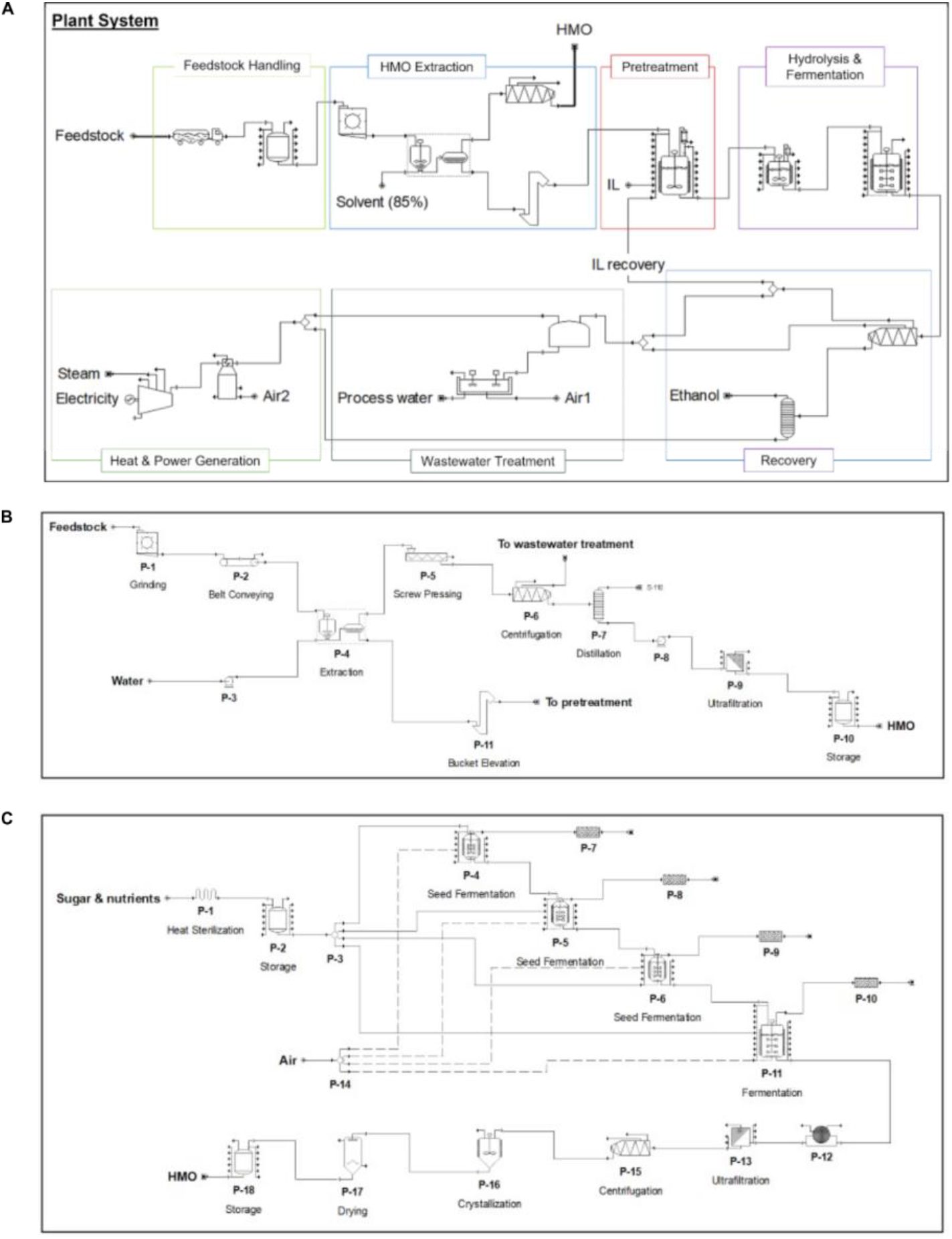
Process models used in this study. A) Process model of cellulosic biorefinery co-producing biofuel and HMOs. B) Process model of LNFPI extraction process in biomass sorghum. C) Process model of HMO production process in *E. coli*.

### Supplementary Tables

**Supplementary Table 1.**
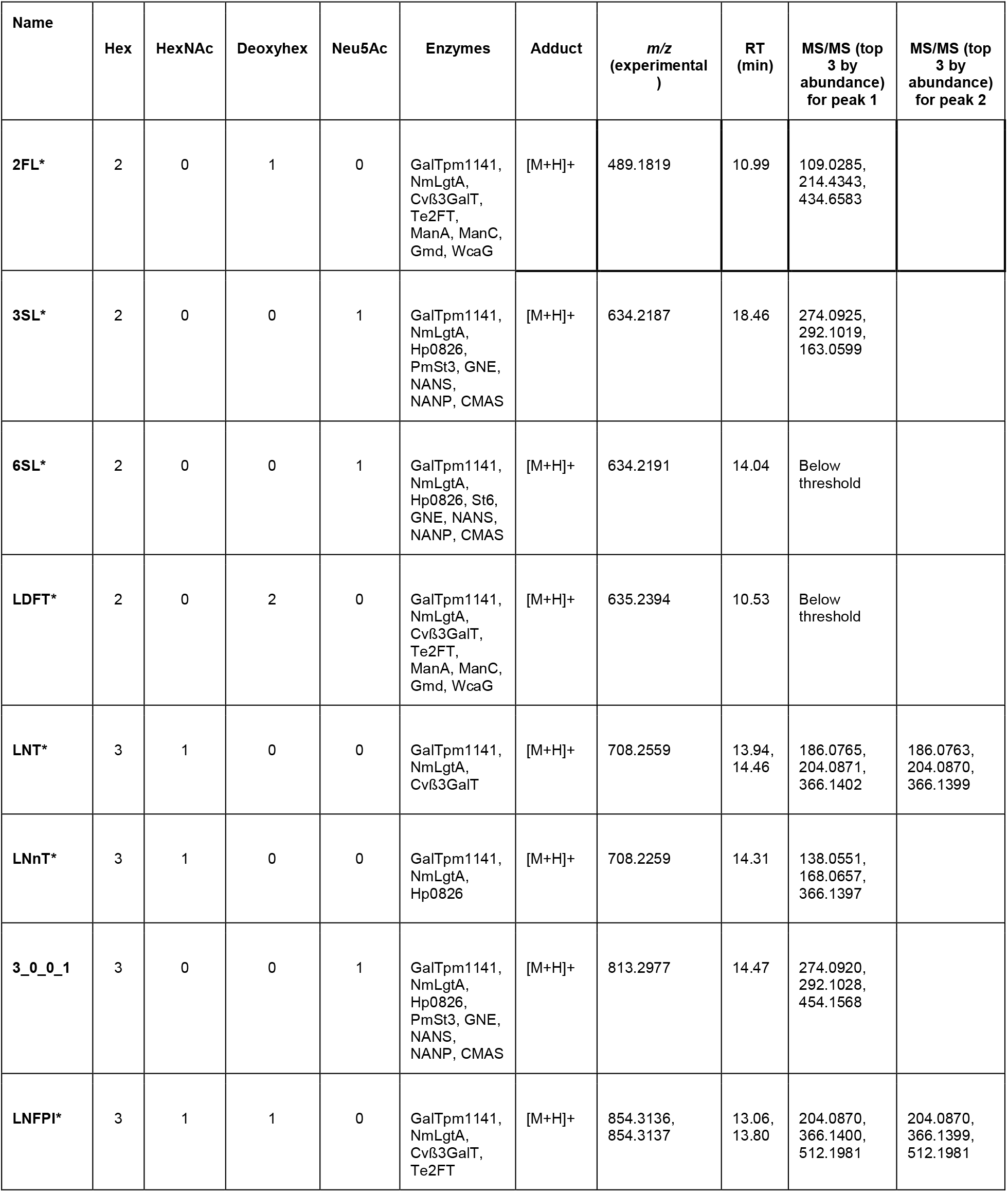
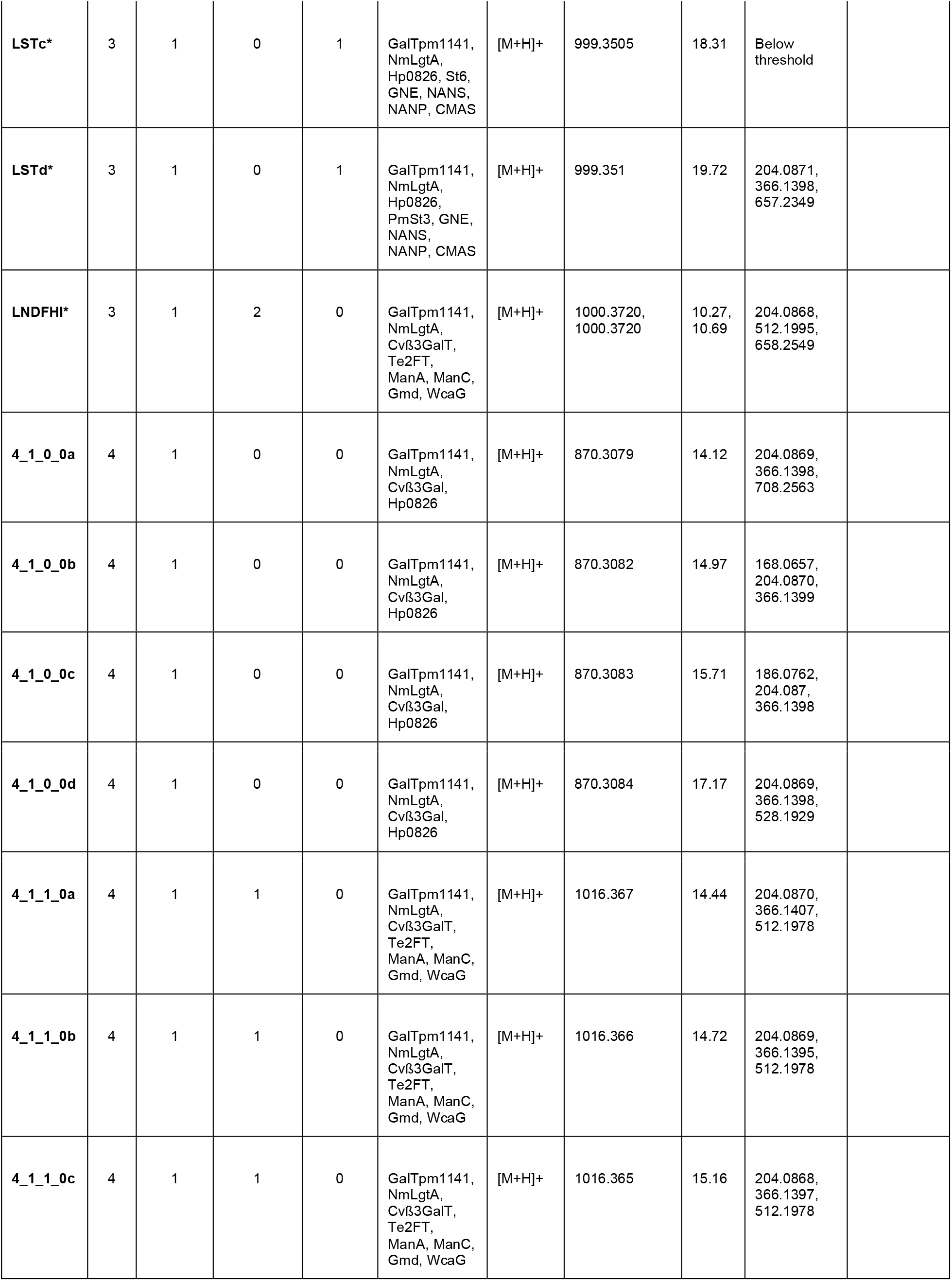
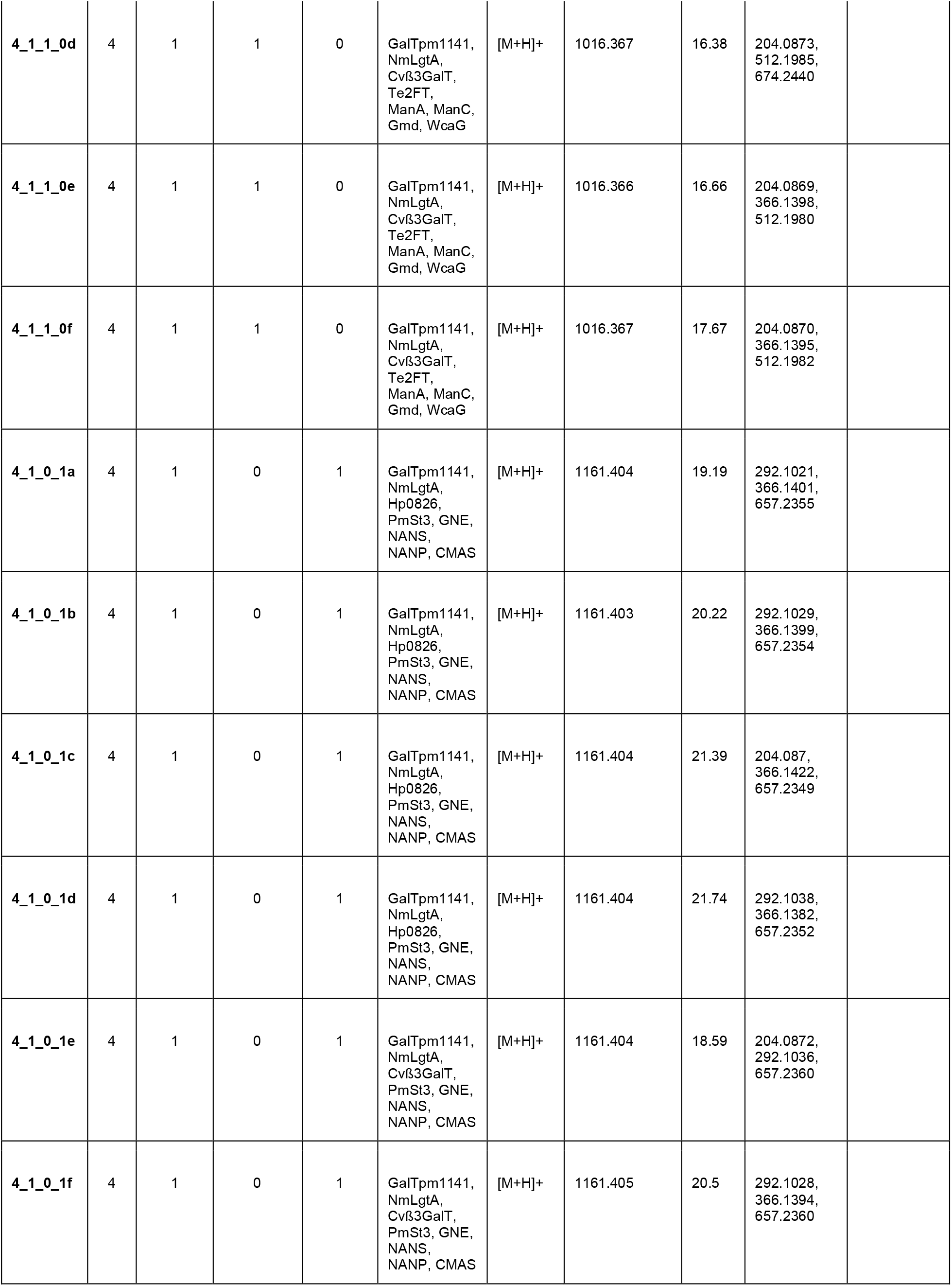
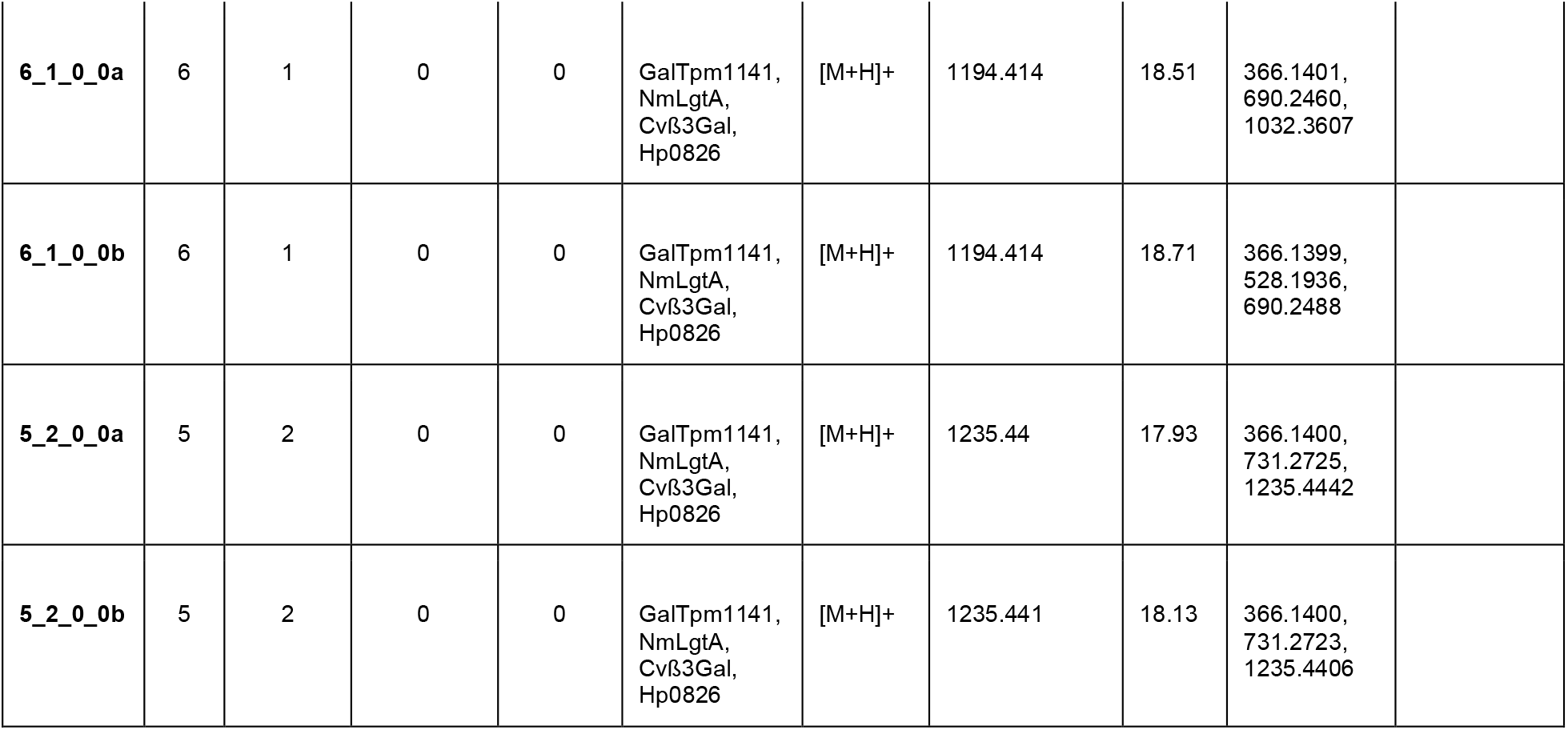
Identification of HMOs through liquid chromatography-mass spectrometry. Obtained with a Thermo Fisher Scientific Q-Exactive mass spetrometer *denotes HMOs identified with use of authenticated standards.

**Supplementary Table 2:**
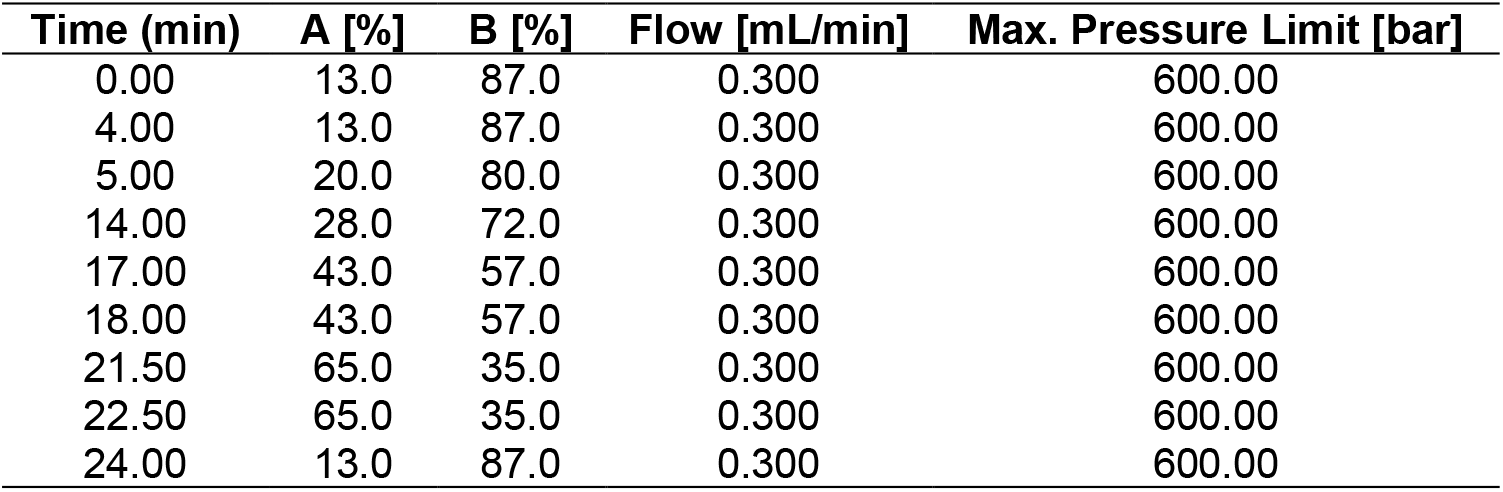
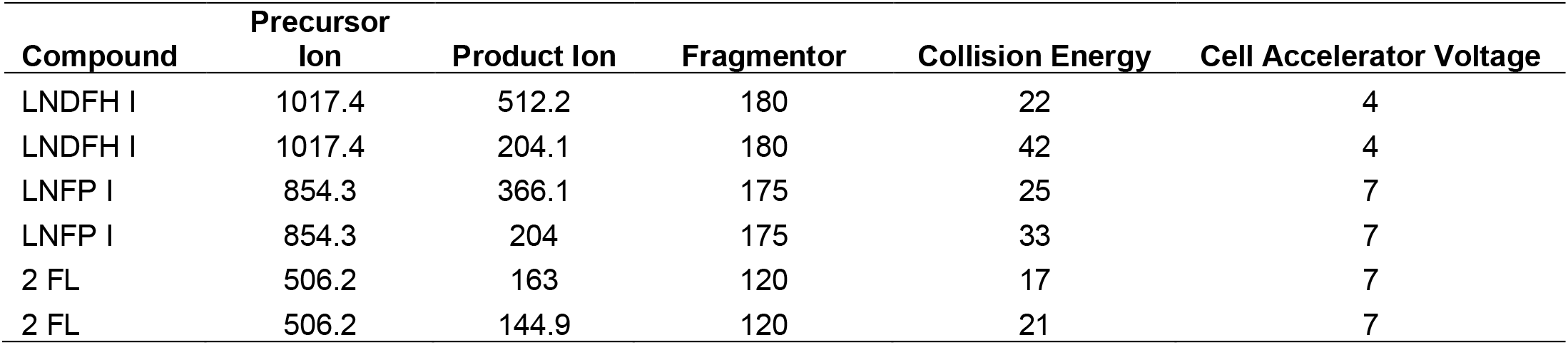
Gradient of the mobile phase for Triple Quadrupole Liquid Chromatography Mass Spectrometry System (QqQ LC-MS) and MRM transitions for HMOs analysis.

| Time (min) | A [%] | B [%] | Flow [mL/min] | Max. Pressure Limit [bar] |
| --- | --- | --- | --- | --- |
| 0.00 | 13.0 | 87.0 | 0.300 | 600.00 |
| 4.00 | 13.0 | 87.0 | 0.300 | 600.00 |
| 5.00 | 20.0 | 80.0 | 0.300 | 600.00 |
| 14.00 | 28.0 | 72.0 | 0.300 | 600.00 |
| 17.00 | 43.0 | 57.0 | 0.300 | 600.00 |
| 18.00 | 43.0 | 57.0 | 0.300 | 600.00 |
| 21.50 | 65.0 | 35.0 | 0.300 | 600.00 |
| 22.50 | 65.0 | 35.0 | 0.300 | 600.00 |
| 24.00 | 13.0 | 87.0 | 0.300 | 600.00 |

| Compound | Precursor Ion | Product Ion | Fragmentor | Collision Energy | Cell Accelerator Voltage |
| --- | --- | --- | --- | --- | --- |
| LNDFH I | 1017.4 | 512.2 | 180 | 22 | 4 |
| LNDFH I | 1017.4 | 204.1 | 180 | 42 | 4 |
| LNFP I | 854.3 | 366.1 | 175 | 25 | 7 |
| LNFP I | 854.3 | 204 | 175 | 33 | 7 |
| 2 FL | 506.2 | 163 | 120 | 17 | 7 |
| 2 FL | 506.2 | 144.9 | 120 | 21 | 7 |

**Supplementary Table 3.** Concentrations of simple sugars and phenolic compounds following purification of HMOs from plant leaves for use in bacterial growth studies. Measurements are in mg/g dry weight (dwt) unless otherwise specified. SD indicates standard deviation. Phenolics are measured as mg/g Gallic acid equivalents.

| Sample | Glucose | Sucrose | Fuctose | Total | Phenolics |
| --- | --- | --- | --- | --- | --- |
| LNFPI+GDP-fucose pathway (PVPP+C18) for microbial growth | 0.038 ± 0.0028 | 0.00092 ± 0.000076 | 0.014 ± 0.0013 | 0.0533 | 0.98 ± 0.0054 |

**Supplementary Table 4.**
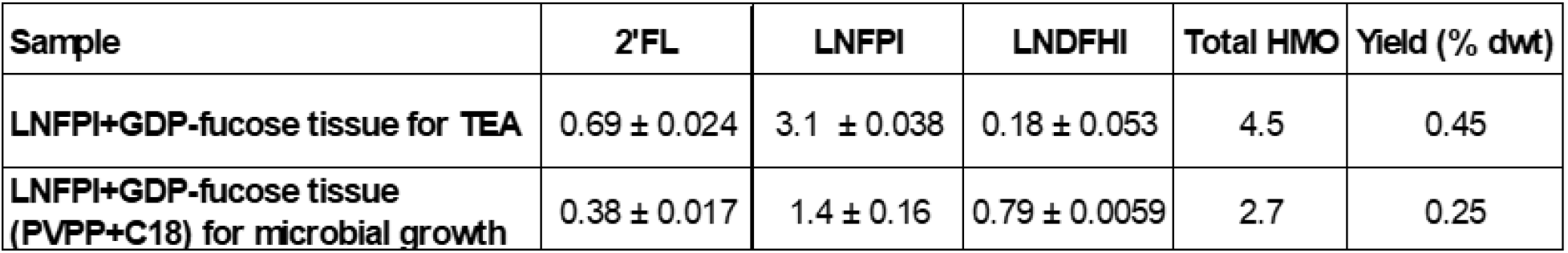
HMO yields following purification for bacterial growth. HMOs were purified from *N. benthamina* transiently expressing the LNFPI and GDP-fucose biosynthetic pathway. Measurements are in mg/g dry weight (dwt) unless otherwise specified. SD indicates standard deviation.

## References

1. Walsh, C., Lane, J. A., van Sinderen, D. & Hickey, R. M. Human milk oligosaccharides: Shaping the infant gut microbiota and supporting health. Journal of Functional Foods 72, 104074 (2020).

2. Kirmiz, N., Robinson, R. C., Shah, I. M., Barile, D. & Mills, D. A. Milk Glycans and Their Interaction with the Infant-Gut Microbiota. Annu Rev Food Sci Technol 9, 429–450 (2018).

3. Vandenplas, Y. et al. Human Milk Oligosaccharides: 2′-Fucosyllactose (2′-FL) and Lacto-N-Neotetraose (LNnT) in Infant Formula. Nutrients 10, 1161 (2018).

4. Wiciński, M., Sawicka, E., Gębalski, J., Kubiak, K. & Malinowski, B. Human Milk Oligosaccharides: Health Benefits, Potential Applications in Infant Formulas, and Pharmacology. Nutrients 12, 266 (2020).

5. CDC. 2022 Breastfeeding Report Card. Centers for Disease Control and Prevention https://www.cdc.gov/breastfeeding/data/reportcard.htm (2022).

6. Lee, S. et al. Human milk oligosaccharide 2’-fucosyllactose supplementation improves gut barrier function and signaling in the vagal afferent pathway in mice. Food Funct 12, 8507– 8521 (2021).

7. Šuligoj, T. et al. Effects of Human Milk Oligosaccharides on the Adult Gut Microbiota and Barrier Function. Nutrients 12, 2808 (2020).

8. Elison, E. et al. Oral supplementation of healthy adults with 2′-O-fucosyllactose and lacto-N-neotetraose is well tolerated and shifts the intestinal microbiota. British Journal of Nutrition 116, 1356–1368 (2016).

9. Iribarren, C. et al. Human milk oligosaccharide supplementation in irritable bowel syndrome patients: A parallel, randomized, double-blind, placebo-controlled study. Neurogastroenterology & Motility 32, e13920 (2020).

10. Iribarren, C. et al. The Effects of Human Milk Oligosaccharides on Gut Microbiota, Metabolite Profiles and Host Mucosal Response in Patients with Irritable Bowel Syndrome. Nutrients 13, 3836 (2021).

11. Bych, K. et al. Production of HMOs using microbial hosts — from cell engineering to large scale production. Current Opinion in Biotechnology 56, 130–137 (2019).

12. Palur, D. S. K., Pressley, S. R. & Atsumi, S. Microbial Production of Human Milk Oligosaccharides. Molecules 28, 1491 (2023).

13. Shani, G. et al. Fucosylated Human Milk Oligosaccharide Foraging within the Species Bifidobacterium pseudocatenulatum Is Driven by Glycosyl Hydrolase Content and Specificity. Appl Environ Microbiol 88, e0170721 (2022).

14. Soyyılmaz, B. et al. The Mean of Milk: A Review of Human Milk Oligosaccharide Concentrations throughout Lactation. Nutrients 13, 2737 (2021).

15. Chen, X. Chapter Four - Human Milk Oligosaccharides (HMOS): Structure, Function, and Enzyme-Catalyzed Synthesis. in Advances in Carbohydrate Chemistry and Biochemistry (eds. Baker, D. C. & Horton, D.) vol. 72 113–190 (Academic Press, 2015).

16. Bar-Peled, M. & O’Neill, M. A. Plant Nucleotide Sugar Formation, Interconversion, and Salvage by Sugar Recycling*. Annual Review of Plant Biology 62, 127–155 (2011).

17. Zhu, F., Du, B. & Xu, B. A critical review on production and industrial applications of beta-glucans. Food Hydrocolloids 52, 275–288 (2016).

18. Palaniappan, A., Antony, U. & Emmambux, M. N. Current status of xylooligosaccharides: Production, characterization, health benefits and food application. Trends in Food Science & Technology 111, 506–519 (2021).

19. Choct, M., Dersjant-Li, Y., McLeish, J. & Peisker, M. Soy Oligosaccharides and Soluble Non-starch Polysaccharides: A Review of Digestion, Nutritive and Anti-nutritive Effects in Pigs and Poultry. Asian-Australasian Journal of Animal Sciences 23, 1386–1398 (2010).

20. Shoaib, M. et al. Inulin: Properties, health benefits and food applications. Carbohydrate Polymers 147, 444–454 (2016).

21. Barnum, C. R., Endelman, B. J. & Shih, P. M. Utilizing Plant Synthetic Biology to Improve Human Health and Wellness. Frontiers in Plant Science 12, 1824 (2021).

22. Kellman, B. P. et al. Elucidating Human Milk Oligosaccharide biosynthetic genes through network-based multi-omics integration. Nat Commun 13, 2455 (2022).

23. Thompson, M. G. et al. Agrobacterium tumefaciens: A Bacterium Primed for Synthetic Biology. BioDesign Research vol. 2020 https://spj.sciencemag.org/journals/bdr/2020/8189219/ (2020).

24. Parschat, K., Schreiber, S., Wartenberg, D., Engels, B. & Jennewein, S. High-Titer De Novo Biosynthesis of the Predominant Human Milk Oligosaccharide 2′-Fucosyllactose from Sucrose in Escherichia coli. ACS Synth. Biol. 9, 2784–2796 (2020).

25. Ooi, K.-E., Zhang, X.-W., Kuo, C.-Y., Liu, Y.-J. & Yu, C.-C. Chemoenzymatic Synthesis of Asymmetrically Branched Human Milk Oligosaccharide Lacto-N-Hexaose. Frontiers in Chemistry 10, (2022).

26. McArthur, J. B., Yu, H. & Chen, X. A Bacterial β1–3-Galactosyltransferase Enables Multigram-Scale Synthesis of Human Milk Lacto-N-tetraose (LNT) and Its Fucosides. ACS Catal. 9, 10721–10726 (2019).

27. Li, Y. et al. Donor substrate promiscuity of bacterial β1–3-N-acetylglucosaminyltransferases and acceptor substrate flexibility of β1–4-galactosyltransferases. Bioorg Med Chem 24, 1696–1705 (2016).

28. Thurl, S., Munzert, M., Boehm, G., Matthews, C. & Stahl, B. Systematic review of the concentrations of oligosaccharides in human milk. Nutr Rev 75, 920–933 (2017).

29. Chen, X., Zhao, C. & Yu, H. Te2ft enzyme for enzymatic synthesis of alpha1-2-fucosides. (2017).

30. Hobbs, M., Jahan, M., Ghorashi, S. A. & Wang, B. Current Perspective of Sialylated Milk Oligosaccharides in Mammalian Milk: Implications for Brain and Gut Health of Newborns. Foods 10, 473 (2021).

31. Castilho, A. et al. Construction of a Functional CMP-Sialic Acid Biosynthesis Pathway in Arabidopsis. Plant Physiol 147, 331–339 (2008).

32. Drouillard, S., Mine, T., Kajiwara, H., Yamamoto, T. & Samain, E. Efficient synthesis of 6′-sialyllactose, 6,6′-disialyllactose, and 6′-KDO-lactose by metabolically engineered E. coli expressing a multifunctional sialyltransferase from the Photobacterium sp. JT-ISH-224. Carbohydrate Research 345, 1394–1399 (2010).

33. Chen, X., Thon, V. & Yu, H. PmST3 enzyme for chemoenzymatic synthesis of alpha-2-3-sialosides. (2017).

34. Ryan, M. d. & Drew, J. Foot-and-mouth disease virus 2A oligopeptide mediated cleavage of an artificial polyprotein. The EMBO Journal 13, 928–933 (1994).

35. Ghoddusi, H. B., Grandison, M. A., Grandison, A. S. & Tuohy, K. M. In vitro study on gas generation and prebiotic effects of some carbohydrates and their mixtures. Anaerobe 13, 193–199 (2007).

36. Sela, D. A. et al. The genome sequence of Bifidobacterium longum subsp. infantis reveals adaptations for milk utilization within the infant microbiome. Proceedings of the National Academy of Sciences 105, 18964–18969 (2008).

37. Underwood, M. A. et al. A Comparison of Two Probiotic Strains of Bifidobacteria in Premature Infants. The Journal of Pediatrics 163, 1585–1591.e9 (2013).

38. Yang, M. et al. Accumulation of high-value bioproducts in planta can improve the economics of advanced biofuels. PNAS 117, 8639–8648 (2020).

39. Yang, M. et al. Comparing in planta accumulation with microbial routes to set targets for a cost-competitive bioeconomy. Proceedings of the National Academy of Sciences 119, e2122309119 (2022).

40. Derya, S. M. et al. Biotechnologically produced fucosylated oligosaccharides inhibit the binding of human noroviruses to their natural receptors. Journal of Biotechnology 318, 31– 38 (2020).

41. Sandalow, D., Aines, R., Friedmann, J., McCormick, C. & Sanchez, D. Biomass Carbon Removal and Storage (BiRCS) Roadmap. LLNL-TR--815200, 1763937, 1024342 https://www.osti.gov/servlets/purl/1763937/<x> (2021) doi:10.2172/1763937.

42. Barnum, C. R., Endelman, B. J., Ornelas, I. J., Pignolet, R. M. & Shih, P. M. Optimization of Heterologous Glucoraphanin Production In Planta. ACS Synth. Biol. 11, 1865–1873 (2022).

43. Engler, C., Gruetzner, R., Kandzia, R. & Marillonnet, S. Golden Gate Shuffling: A One-Pot DNA Shuffling Method Based on Type IIs Restriction Enzymes. PLOS ONE 4, e5553 (2009).

44. Gibson, D. G. et al. Enzymatic assembly of DNA molecules up to several hundred kilobases. Nat Methods 6, 343–345 (2009).

45. Froger, A. & Hall, J. E. Transformation of plasmid DNA into E. coli using the heat shock method. J Vis Exp 253 (2007) doi:10.3791/253.

46. Lin, J.-J. Electrotransformation of Agrobacterium. in Electroporation Protocols for Microorganisms (ed. Nickoloff, J. A.) 171–178 (Humana Press, 1995). doi:10.1385/0-89603-310-4:171.

47. Lakatos, L., Szittya, G., Silhavy, D. & Burgyán, J. Molecular mechanism of RNA silencing suppression mediated by p19 protein of tombusviruses. The EMBO Journal 23, 876–884 (2004).

48. Xu, G. et al. Absolute Quantitation of Human Milk Oligosaccharides Reveals Phenotypic Variations during Lactation. The Journal of Nutrition 147, 117–124 (2017).

49. Tsugawa, H. et al. MS-DIAL: data-independent MS/MS deconvolution for comprehensive metabolome analysis. Nat Methods 12, 523–526 (2015).

50. de Moura Bell, J. M. L. N., et al. An integrated bioprocess to recover bovine milk oligosaccharides from colostrum whey permeate. Journal of Food Engineering 216, 27–35 (2018).

51. Magalhães, P. J. et al. Isolation of phenolic compounds from hop extracts using polyvinylpolypyrrolidone: Characterization by high-performance liquid chromatography– diode array detection–electrospray tandem mass spectrometry. Journal of Chromatography A 1217, 3258–3268 (2010).

52. Laurentin, A. & Edwards, C. A. A microtiter modification of the anthrone-sulfuric acid colorimetric assay for glucose-based carbohydrates. Analytical Biochemistry 315, 143–145 (2003).

53. Singleton, V. L., Orthofer, R. & Lamuela-Raventós, R. M. [14] Analysis of total phenols and other oxidation substrates and antioxidants by means of folin-ciocalteu reagent. in Methods in Enzymology vol. 299 152–178 (Academic Press, 1999).

54. Huang, Y.-P., Paviani, B., Fukagawa, N. K., Phillips, K. M. & Barile, D. Comprehensive oligosaccharide profiling of commercial almond milk, soy milk, and soy flour. Food Chemistry 409, 135267 (2023).

55. Huang, Y.-P., Robinson, R. C. & Barile, D. Food glycomics: Dealing with unexpected degradation of oligosaccharides during sample preparation and analysis. Journal of Food and Drug Analysis 30, 62–76 (2022).

56. Bhattacharya, M., Salcedo, J., Robinson, R. C., Henrick, B. M. & Barile, D. Peptidomic and glycomic profiling of commercial dairy products: identification, quantification and potential bioactivities. NPJ Sci Food 3, 4 (2019).

57. Lee, H., Garrido, D., Mills, D. A. & Barile, D. Hydrolysis of milk gangliosides by infant-gut associated bifidobacteria determined by microfluidic chips and high-resolution mass spectrometry. ELECTROPHORESIS 35, 1742–1750 (2014).

58. Humbird, D., et al. Process Design and Economics for Biochemical Conversion of Lignocellulosic Biomass to Ethanol: Dilute-Acid Pretreatment and Enzymatic Hydrolysis of Corn Stover. NREL/TP-5100-47764, 1013269 http://www.osti.gov/servlets/purl/1013269/<x> (2011) doi:10.2172/1013269.

